# Chemical precision glyco-mutagenesis by glycosyltransferase engineering in living cells

**DOI:** 10.1101/669861

**Authors:** Benjamin Schumann, Stacy A. Malaker, Simon P. Wisnovsky, Marjoke F. Debets, Anthony J. Agbay, Daniel Fernandez, Lauren J. S. Wagner, Liang Lin, Junwon Choi, Douglas M. Fox, Jessie Peh, Melissa A. Gray, Kayvon Pedram, Jennifer J. Kohler, Milan Mrksich, Carolyn R. Bertozzi

## Abstract

Studying posttranslational modifications classically relies on experimental strategies that oversimplify the complex biosynthetic machineries of living cells. Protein glycosylation contributes to essential biological processes, but correlating glycan structure, underlying protein and disease-relevant biosynthetic regulation is currently elusive. Here, we engineer living cells to tag glycans with editable chemical functionalities while providing information on biosynthesis, physiological context and glycan fine structure. We introduce a non-natural substrate biosynthetic pathway and use engineered glycosyltransferases to incorporate chemically tagged sugars into the cell surface glycome of the living cell. We apply the strategy to a particularly redundant yet disease-relevant human glycosyltransferase family, the polypeptide *N*-acetylgalactosaminyl transferases. This approach bestows a gain-of-function modification on cells where the products of individual glycosyltransferases can be selectively characterized or manipulated at will.

## Main Text

Posttranslational modifications expand the structural diversity of proteins, but are notoriously difficult to study in living systems. Most modifications are refractory to direct genetic manipulation and require reductionist strategies such as *in vitro* systems or simplified cells. Glycans are the prime example for this: the human glycome is constructed by the combinatorial activity of more than 250 glycosyltransferases (GTs) with both hierarchical and competing activities. On the cell surface, glycans play a central role in modulating signal transduction, cell-cell interactions and biophysical properties of the plasma membrane (*1, 2*). Understanding and manipulating these processes should thus be possible by adorning specific glycans with editable chemical functionalities. In such a synthetic biology approach, individual GTs could be engineered to accommodate a chemical functionality that is not found in native substrates and not accommodated by other GTs. This “bump-and-hole” tactic has been applied to a range of enzymes by us and others (*3–9*), but application in the living cell is a substantial technical challenge: the nucleotide-based substrate analog must be delivered across the plasma membrane and the cell must stably express the correctly localized and folded mutant enzyme. Bump-and-hole engineering is particularly attractive to deconvolve GT families of multiple homologous isoenzymes: the complex interplay of these isoenzymes in the secretory pathway cannot be probed in sufficient detail in *in vitro* assays.

One of the largest GT families in the human genome is the polypeptide *N*-acetylgalacosaminyl (GalNAc) transferase family (GalNAc-T-1…T-20). Transferring GalNAc to Ser/Thr side chains, GalNAcTs initiate abundant O-linked glycosylation in the secretory pathway (Fig. 1A) (*10, 11*). O-GalNAc glycosylation can differ based on cell type or activation stage, and clear disease phenotypes are associated with glycan aberration (*12, 13*). Unsurprisingly, GalNAcT expression is often associated with tumorigenesis, sometimes with opposite effects on different types of cancer (*14, 15*). However, the absence of a glycosylation consensus sequence and the variability of glycan elaboration render O-GalNAc glycans challenging to study by mass spectrometry (MS) based glycoproteomics. Thus, the protein substrate specificity of each isoenzyme is poorly studied and largely built on data inferred from synthetic peptide libraries (*16*). So-called SimpleCells that do not elaborate O-GalNAc glycans make glycoproteins that are easier to enrich and study by MS (*17–20*). Knock-outs of single GalNAcTs in the SimpleCell background have been profiled by glycosproteomics (*18, 21*). Similarly, glycopeptides from non-SimpleCells with titratable GalNAcT knock-in have been enzymatically simplified to allow for enrichment and MS (*22*). These studies have revealed comprehensive glycoproteomics datasets, but suffer from loss of glycan elaboration and laborious genome engineering required for targeted knock-in. Further, the activity of GalNAcT isoenzymes is both redundant and competitive, such that compensation and/or shift of glycosylation sites occurs upon knock-out (*21, 22*). A gain-of-function strategy to visualize the products of a particular GT on the living cell is currently elusive.

**Fig. 1:**
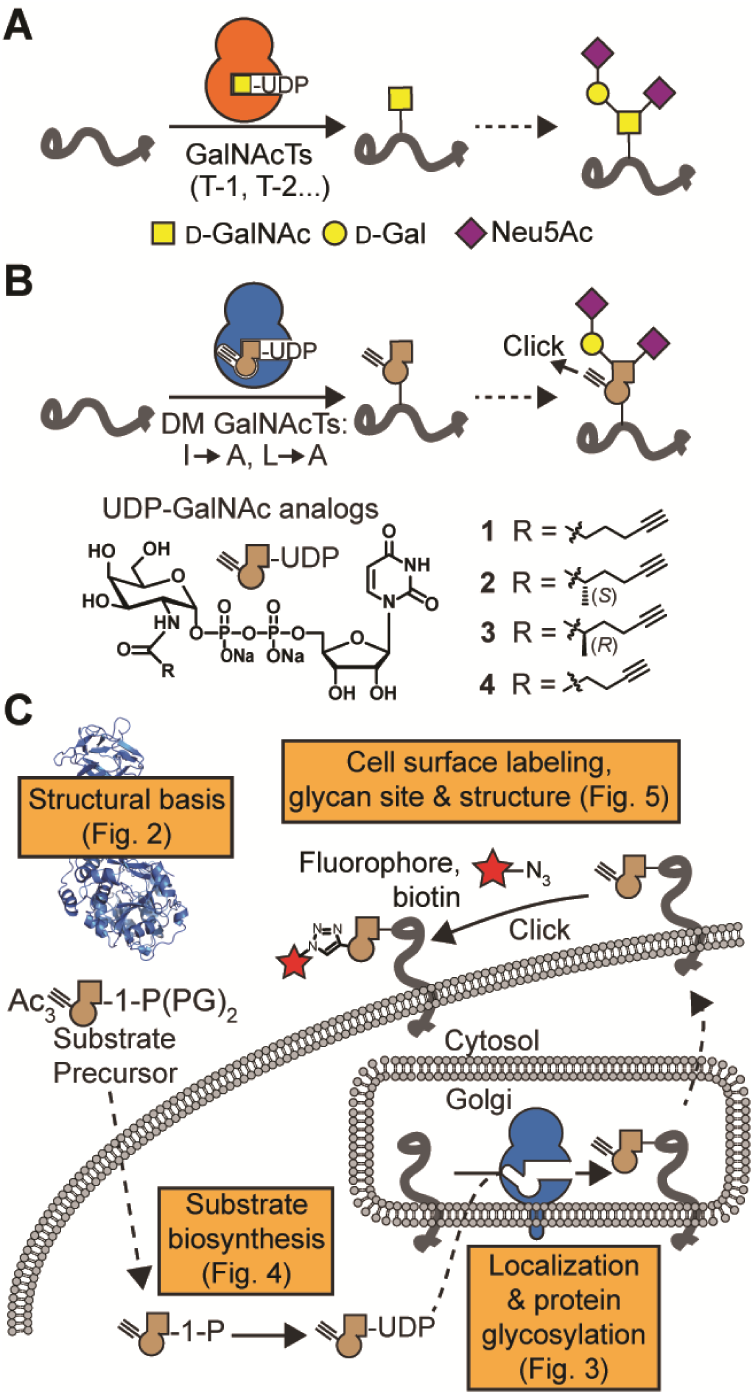
GalNAcT bump-and-hole engineering. *A*, GalNAcTs initiate O-GalNAc glycosylation. Transfer of GalNAc to a Ser or Thr side chain is followed by downstream glycan elongation. *B*, the principle of bump-and-hole engineering. Engineered double mutant (DM) GalNAcTs are paired with UDP-GalNAc analogs **1**-**4** to chemically tag GalNAcT substrates that can be monitored by click chemistry. *C*, Overview of the steps taken in this study toward GalNAcT bump-and-hole engineering in the living cell. PG = Protecting group.

Here, we equipped living cells with the ability to tag the protein substrates of individual GalNAcTs with chemical, editable functionalities. We made use of the fact that a double mutation (“DM”, I253A/L310A double mutant in T-2, fig. 1B) re-programmed GalNAcTs to accept alkyne-containing uridine diphosphate (UDP)-GalNAc analogs such as compounds **1**-**4** instead of the native substrate UDP-GalNAc (Fig. 1B) (*8*). We endowed living cells with the capacity to biosynthesize a UDP-GalNAc analog that was complementary to engineered DM-GalNAcTs. Further, we showed that DM-GalNAcTs emulate wild type (WT) GalNAcTs with regard to structure, localization and protein substrate specificity. This approach bestowed a bioorthogonal tag (*23–27*) on the protein substrates of distinct GalNAcT isoenzymes while conserving the complexity of glycan elaboration in the secretory pathway (Fig. 1C). Precision glycome engineering has widespread applications in biomarker discovery, GT profiling and targeted cell surface engineering.

### Structural basis for GalNAcT engineering

As bump-and-hole engineering of a GT family in living cells has no precedent, we first set out to understand the structural implications of this approach. All GalNAcTs are type II transmembrane proteins with luminal GT and lectin domains connected through a flexible linker (*10, 11, 28*). We crystallized the luminal part of DM-T-2 in complex with the native ligands Mn^2+^, UDP, and the substrate peptide EA2 (PTTDSTTPAPTTK) at 1.8 Å resolution. Comparison of DM-T-2 and WT-T-2 (PDB 2FFU) revealed complete conservation of both the three-dimensional enzyme architecture and bound ligand structure (Fig. 2A, fig. S1 and Table S1) (*29*). In DM-T-2, the interdomain linker adopts an extended conformation previously found in the catalytically active WT enzyme (*30, 31*). The mutant A253 and A310 side chains in the DM enzyme are congruent with the WT side chains I253 and L310, respectively (Fig. 2B). Two glycine residues, G308 and G309, are slightly shifted by 1.2 Å and 1.7 Å (Cα distances), respectively, likely to account for the changes elsewhere in the active site. DM-T-2 thus retains the native structural properties of the WT-GalNAcT enzyme.

**Fig. 2:**
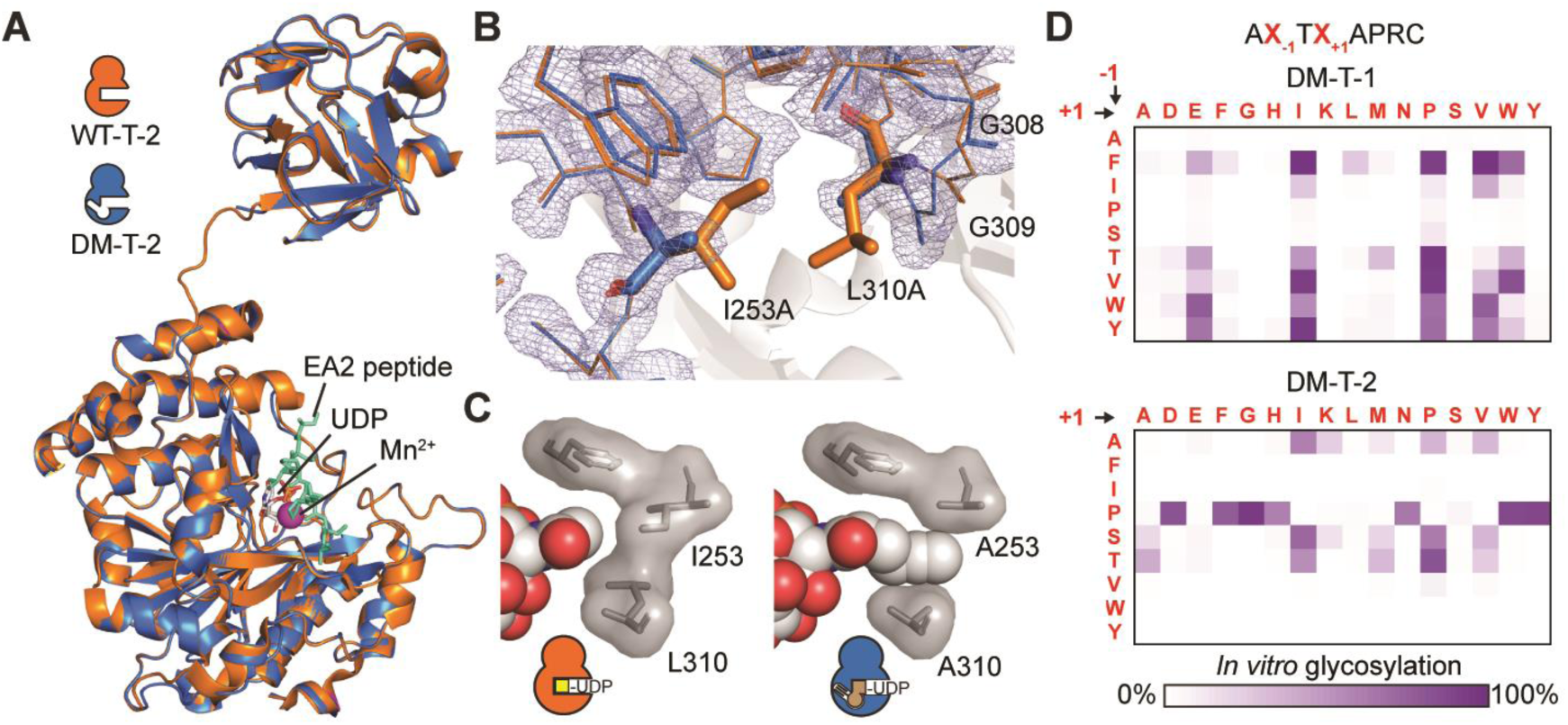
Bump-and-hole engineering conserves folding and substrate binding of GalNAcT-2. *A*, crystal structure of DM-T-2 at 1.8 Å superposed with WT-T-2 (PDB 2FFU). Bound EA2 substrate peptide is cyan (sticks), Mn^2+^ magenta (sphere) and UDP grey (sticks). Ligands are taken from DM-T-2. For superposition with WT-T-2 ligands, see figure S1A. *B*, superposition of the UDP-sugar binding site of DM-T-2 and WT-T-2. Electron density is rendered at 1 σ and carved at 1.6 Å. *C*, depiction of UDP-GalNAc analog **1** in a co-crystal structure with DM-T-2 at 3.1 Å, and UDP-GalNAc in a co-crystal structure with WT-T-2 (PDB 4D0T) (*30*), as well as WT and mutated gatekeeper residues. *D*, substrate specificities of DM-T1 and DM-T-2 as determined in an *in vitro* glycosylation assay with detection by SAMDI-MS. For comparison with WT-GalNAcT glycosylation, see figure S2. Data are from one representative out of two independent experiments. A co-crystal structure of DM-T-2, Mn^2+^ and UDP-GalNAc analog **1** at 3.1 Å resolution helped us visualize how the DM-T-2 active site mutations affect enzyme-substrate binding. In comparison with a corresponding WT-T-2/UDP-GalNAc/Mn^2+^/EA2 complex (PDB 4D0T), UDP-sugar binding is completely conserved (Fig. S1B, C and table S1) (*30*). DM-T-2 indeed contains a hole that accommodates the alkyne side chain bump in UDP-GalNAc analog **1** (Fig. 2C). The formation of additional van der Waals interactions between enzyme and substrate explain why the KM of DM-T-2 toward **1** is approximately ten-fold lower than the KM of WT-T-2 toward UDP-GalNAc (*8*).

To corroborate our structural interpretation that GalNAcT bump-and-hole engineering does not alter substrate peptide binding, we profiled the substrate specificities of two engineered GalNAcTs, DM-T-1 and DM-T-2 (*8, 33-35*). A peptide library containing a single Thr residue with randomized neighboring residues was used in *in vitro* glycosylation experiments and analyzed by self-assembled monolayers for matrix-assisted desorption/ionization mass spectrometry (SAMDI-MS, fig. 2D and fig. S2) (*32*). We found a striking difference in substrate specificity between DM-T-1 and DM-T-2 that was similar to the differences seen in the corresponding WT-GalNAcTs: T-1 generally preferred hydrophobic amino acids at -1-position and Glu, Ile, Pro, Val or Trp at +1-position of the Thr glycan acceptor, whereas T-2 showed a preference for Pro, Ala, Ser and Thr at-1-position (Fig. S2) (*32*). These data indicate that bump-and-hole engineering faithfully reports on GalNAcT isoenzyme activity, a crucial prerequisite for a cellular bump-and-hole system.

### DM-GalNAcTs glycosylate membrane proteins

We next sought to confirm that DM-GalNAcTs localize to the Golgi compartment and glycosylate proteins in a membrane environment. Full-length WT- and DM-T-1 and -T-2 with a C-terminal VSV-G epitope tag exhibited doxycycline (Dox) dose-dependent expression under the control of a tetracycline-responsive promoter in stably transfected HepG2 cells (Fig. 3A and fig. S3A) (*36*). All tested GalNAcTs co-localized with the Golgi marker giantin, confirming native localization of engineered GalNAcTs (Fig. 3B and fig. S3B). To assess GalNAcT activity, membrane protein fractions were prepared from WT- and DM-GalNAcT expressing HepG2 cells and used for *in vitro* glycosylation experiments. After incubation with alkyne-containing UDP-GalNAc analog **1**, alkyne-tagged glycoproteins were derivatized with azide-biotin using Cu^I^-catalyzed [3+2] “click” cycloaddition, and characterized on a streptavidin blot. Compounds **1, 2** and **3** were specific substrates for DM-but not WT-GalNAcTs. In the presence of these substrates, DM-T-1 and DM-T-2 labeled overlapping and also unique glycoprotein species (Fig. 3C and fig. S3C, D). In contrast, treatment of lysates from WT-T-1 and -2 expressing cells with compound **4** with a shorter alkyne chain led to substantial labeling, consistent with **4** being a substrate for WT-GalNAcTs (*8, 26*). These results were confirmed using soluble GalNAcTs and a membrane fraction of non-transfected cells (Fig. S3E). Importantly, profiling both enzyme activity and protein specificity of single GalNAcT isoenzymes has been impossible to date even in cell lysates. We next targeted a GalNAcT bump-and-hole platform in the living cell.

**Fig. 3:**
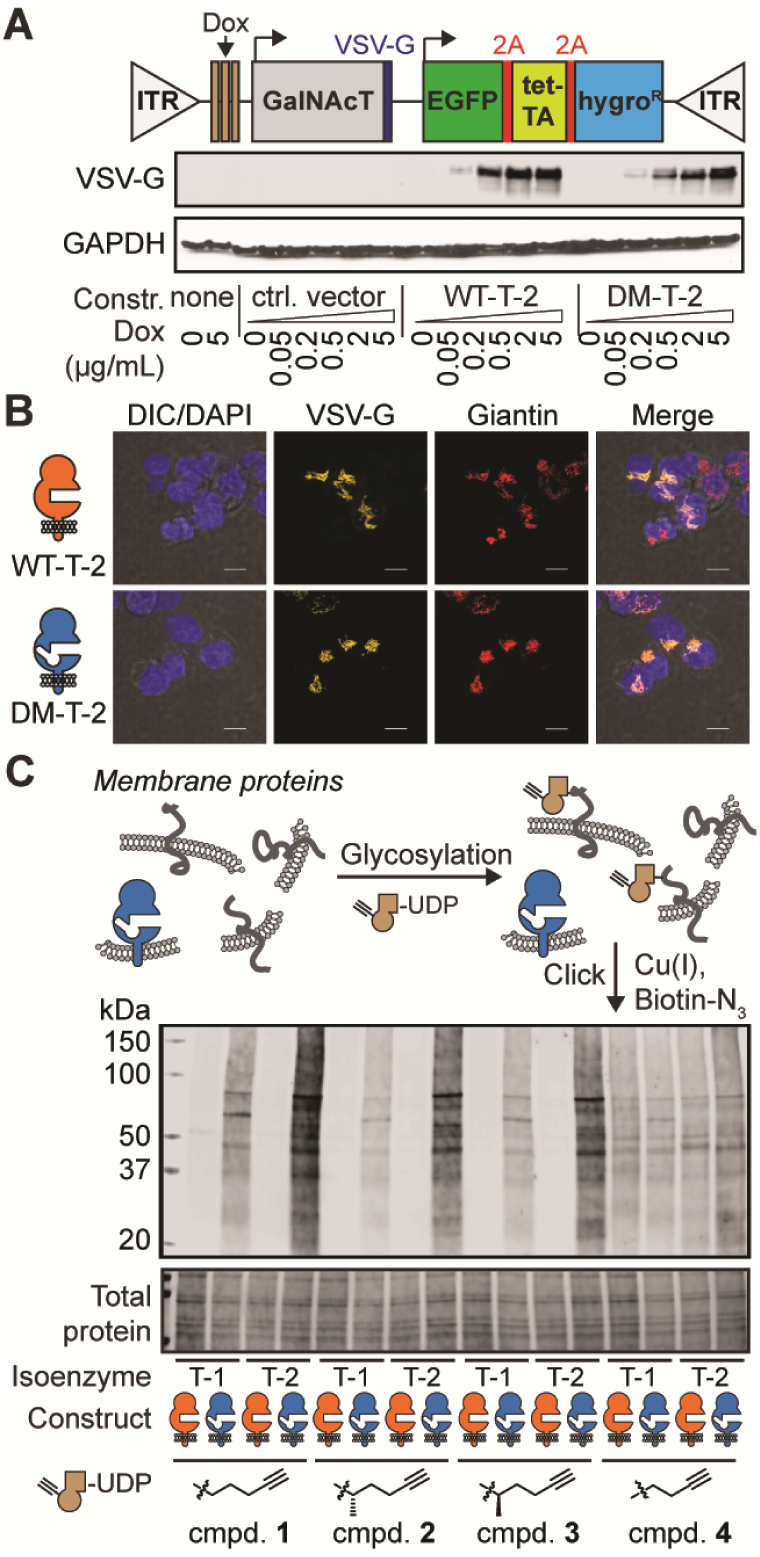
Engineered GalNAcTs localize to the Golgi compartment and glycosylate protein substrates. *A*, expression construct for full-length GalNAcTs under the control of a Dox-inducible promoter. Inverted Tandem Repeats (ITR) are recognized by Sleeping Beauty transposase. WT-T-2 and DM-T-2 were expressed by stably transfected HepG2 cells in a Dox-inducible fashion. *B*, fluorescence microscopy of HepG2 cells stably transfected with T-2 constructs, induced with 0.2 µg/mL Dox and subsequently stained. *C, in vitro* glycosylation of proteins in a membrane fraction by full-length GalNAcTs using UDP-GalNAc analogs. Data are from one representative out of two independent experiments. Experiments were repeated with the membrane fraction of non-transfected cells and soluble, purified GalNAcTs as an enzyme source (Fig. S3). DIC = Differential interference contrast; TA = Trans-activator.

### Biosynthesis of a bumped UDP-GalNAc analog

Among the insightful bump-and-hole studies that have probed various enzyme families, few have been performed in living cells due to the inability to deliver negatively charged substrates across the plasma membrane (*5, 6, 9, 37*). GalNAc analogs have been fed to cells as membrane-permeable per-acetylated precursors that are deprotected by esterases and converted to UDP-GalNAc analogs via the kinase GALK2 and the pyrophosphorylase AGX1 (Fig. 4A) (*26, 27, 38, 39*). By delivery of the corresponding protected GalNAc-1-phosphate analog, a GalNAc analog can bypass GALK2, such that only AGX1 is necessary for biosynthesis to the UDP-GalNAc analog. However, in accordance with previous findings (*39*), bumped UDP-GalNAc analogs **1, 2** and **3** were not biosynthesized from their sugar-1-phosphate precursors in the living cell (Fig. 4A, B).

**Fig. 4:**
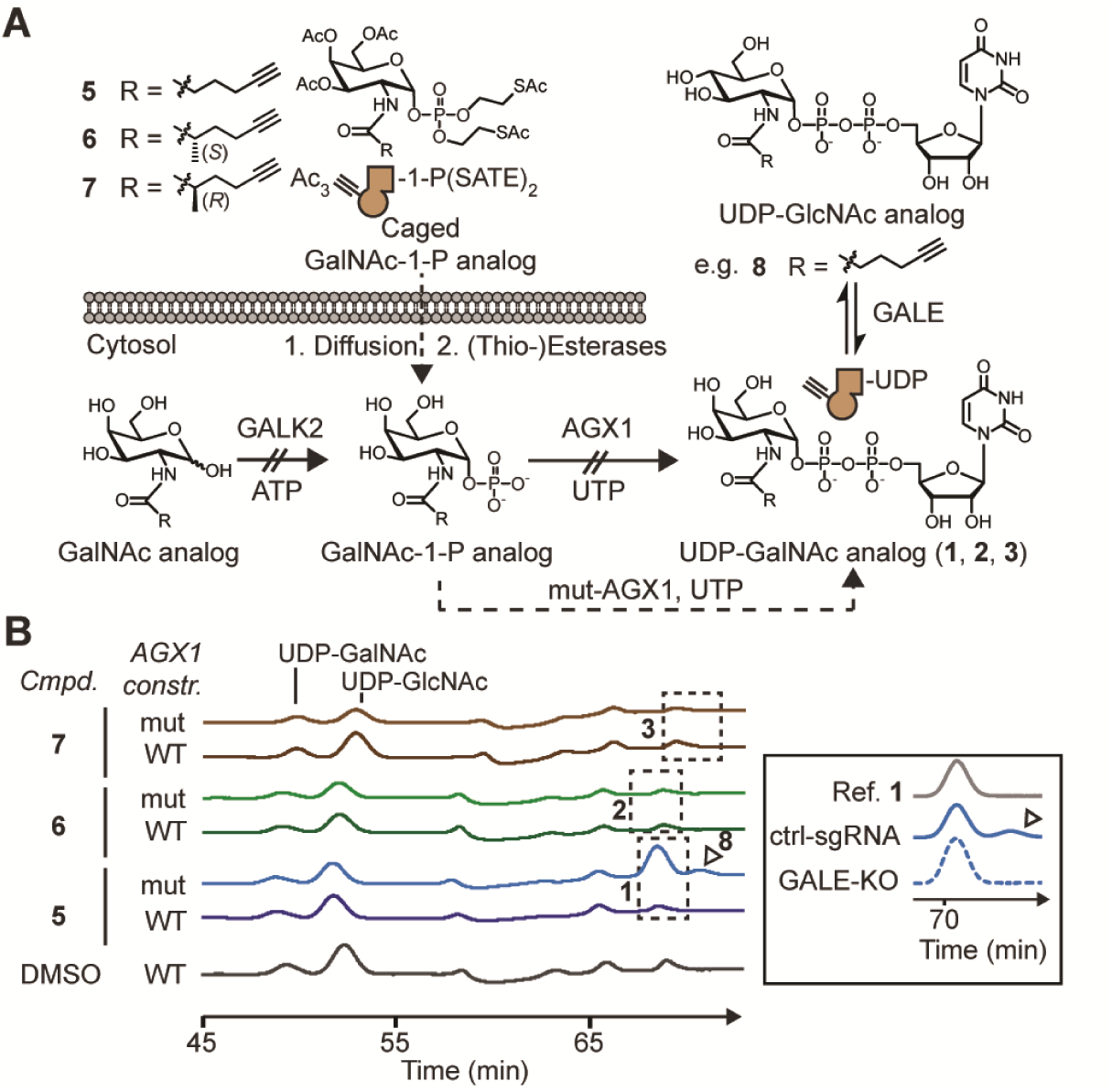
Substrate delivery to the cytosol of living cells. *a*, schematic of substrate delivery. Non-permissive steps are indicated by crossed arrows. The epimerase GALE interconverts UDP-GlcNAc and UDP-GalNAc. *b*, HPAEC-PAD traces of extracts from HEK293T cells stably expressing WT-AGX1 or mut-AGX1 and fed with the indicated compounds. Dashed boxes indicate retention times of standards in separate reference runs (Fig. S4B). The product of potential epimerization of **1** by GALE, compound **8**, is marked with an arrowhead. Data is of one experiment and was repeated for compound **5** in HEK293T cells transiently transfected with AGX1 constructs, as well as stably transfected K-562 cells (Fig. S4B and fig. S5). Insert: epimerization to **8** is suppressed in GALE-deficient K-562 cells expressing mut-AGX1 and fed with **5**, but not cells carrying a control single guide (sg)RNA. A reference trace of compound **1** is shown. Data are of one representative out of two independent experiments.

A lack of AGX1 activity towards synthetic *N*-acetylglucosamine (GlcNAc) analogs has previously prompted the engineering of AGX1 to recognize the corresponding GlcNAc-1-phosphate analog as a substrate (*40*). As WT-AGX1 accepts both GlcNAc-1-phosphate and GalNAc-1-phosphate as substrates, we investigated whether AGX1 mutants could biosynthesize UDP-GalNAc analogs **1, 2** and **3** from the corresponding GalNAc-1-phosphate analogs in the living cell. We mutated the gatekeeper residues F381 and F383 to Gly or Ala in FLAG-tagged AGX1 expression constructs (*40*). GalNAc-1-phosphate analogs were delivered to stably transfected HEK293T cells by virtue of caged precursors **5, 6** and **7**. UDP-sugar biosynthesis was then determined by high performance anion exchange chromatography with pulsed amperometric detection (HPAEC-PAD) of cell lysates. Gly and Ala mutants of F383 efficiently biosynthesized bumped UDP-GalNAc analog **1**, while neither F381 single mutants nor any F381/F383 double mutants produced **1** despite equal expression levels (Fig. 4A) (*40*). In contrast to linear alkyne **1**, neither methylated alkynes (**2** or **3**) were biosynthesized by engineered AGX1^F383A^, hereby called mut-AGX1 (Fig. 5B and fig. S4B). We thus concluded that bumped GalNAc-1-P(SATE)2 precursor **5** can be used in conjunction with mut-AGX1 to deliver UDP-GalNAc analog **1** to the living cell and establish a GalNAcT bump- and-hole system.

**Fig. 5:**
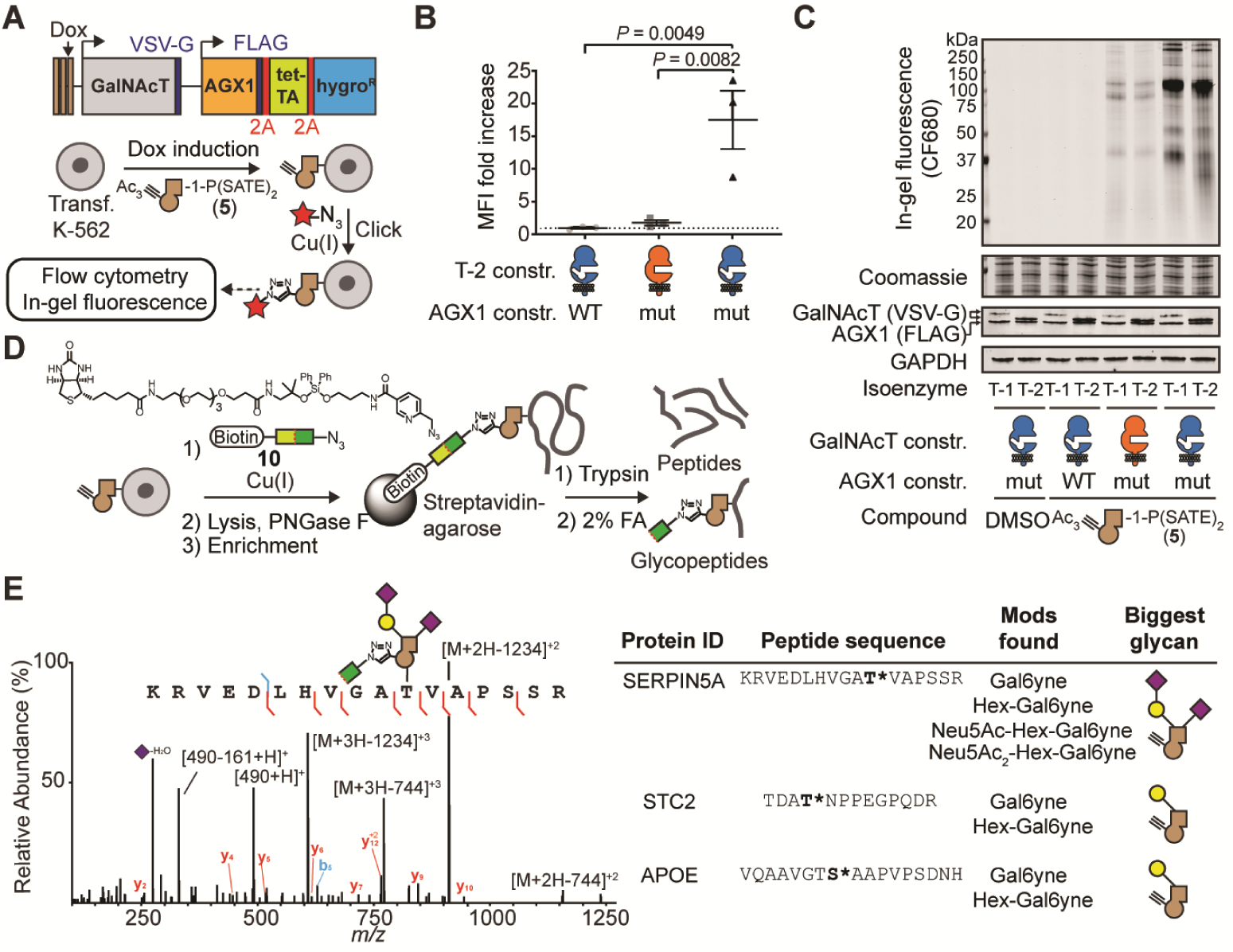
Selective bioorthogonal labeling of the living cell surface with bump-and-hole engineered GalNAcTs. *A*, GalNAcT and AGX1 co-expression construct and workflow of cell surface labeling. Red star depicts a fluorophore. *B*, labeling analysis of K-562 GALE-KO cells by flow cytometry of MB488-picolyl azide labeled and intracellular VSV-G-stained cells. Data are individual values from three independent experiments, means ± SEM of MB488 median fluorescence intensity of VSV-G positive cells. Statistical analysis was performed by two-tailed ratio paired t-test. *C*, labeling analysis by in-gel fluorescence of PNGase F-treated lysates from metabolically labeled K-562 cells. Data is representative of three independent experiments. *D*, schematic of glycoprotein enrichment and on-bead digest. The bifunctional molecule **10** bears an acid-labile diphenyldisiloxane moiety. *E*, exemplary mass spectrometry data: mass spectrum (HCD) of a fully elaborated glycopeptide from SERPIN5A (site Thr39), and further examples from T-2-specific sites from STC2 (Thr28) and APOE (S308). FA = formic acid; MFI = mean fluorescence intensity

In HPAEC-PAD chromatograms of cells biosynthesizing UDP-GalNAc analog **1**, we consistently found a satellite peak that eluted after **1** (arrowhead in figure 4B). The epimerase GALE maintains a cytosolic equilibrium between UDP-GalNAc and UDP-GlcNAc and has been shown to also accept azide-containing analogs (*38, 41*). We found that GALE epimerizes UDP-GalNAc analog **1** to the corresponding UDP-GlcNAc analog **8** (Fig. 4A): first, HPAEC-mass spectrometry confirmed that the satellite peak was caused by an isomer of UDP-GalNAc analog **1** with 328 m/z [M-2H]^2-^. Second, CRISPR-mediated GALE-KO in the K-562 background abrogated the satellite peak (Fig. 4B and fig. S5). In contrast, epimerization was still observed in GALE-containing cells. Alkyne-containing UDP-GlcNAc analog **8** might result in labeling of GlcNAc-containing glycans on the cell surface, such as N-linked glycans. We thus concluded that epimerization of **1** to **8** must be accounted for in glycoproteomics experiments, for example by enzymatic abrogation of cell surface N-glycans during sample processing.

### A glyosyltransferase bump-and-hole-system in living cells

With a GalNAcT bump-and-hole system and a method for cellular substrate delivery of **1** in hand, we set out to probe isoenzyme-dependent glycosylation in the living cell. We cloned FLAG-tagged WT- or mut-AGX1 under a constitutive promoter into an expression vector containing VSV-G-tagged WT or DM versions of GalNAcT-1 or T-2 under the control of a Dox-inducible promoter (Fig. 5A). This setup allowed us to systematically assess any potential background protein labeling when UDP-GalNAc analog **1** could not be biosynthesized, or when **1** was biosynthesized but DM-GalNAcTs were absent. To preclude any labeling due to epimerization of **1** to UDP-GlcNAc analog **8**, we first used GALE-KO K-562 cells for labeling. Cells were supplemented with GalNAc to compensate for the loss of UDP-GalNAc biosynthesis (Fig. S5A). GalNAcT expression was induced (Fig. S6A), cells were fed with caged GalNAc-1-phosphate analog **5**, and the cell surface was reacted with clickable Alexa488-picolyl azide in the presence of a non-membrane-permissive Cu(I) complex (*3, 42*). After gating for positive VSV-G signal, flow cytometry showed a more than 15-fold fluorescence increase when both DM-T-2 and substrate **1** were present over controls lacking either (Fig. 5B and fig. S6B). In GALE-containing cells, a more than 2-fold higher signal was still measured over control cells when a functional bump-and-hole system was present (Fig. S6C).

In order to better visualize the scope of protein labeling by our bump-and-hole system, we reacted cell surfaces with a clickable version of the infrared dye CF680 to profile labeled cell surface glycoproteins by in-gel fluorescence. We compared protein labeling patterns of GALE-containing K-562 cells stably expressing T-1 or T-2 constructs. Here, N-linked glycans were removed by PNGase F treatment prior to analysis to reduce background fluorescence (Fig. S7A). A profound band pattern was observed when functional bump-and-hole pairs (**5**, mut-AGX1 and DM-GalNAcTs) were present (Fig. 5C and fig. S7B). Mut-AGX1 was required for labeling, confirming that our experiments probe enzymatic glycosylation rather than non-specific protein modification (*43*). Fluorescence was of similar intensity as in cells treated with well-characterized alkyne-containing *N*-acetylneuraminic acid precursor Ac4ManNAlk (*44*). Furthermore, the presence of DM-GalNAcT protein was essential, as omission of Dox induction prevented fluorescent labeling (Fig. S7B). Importantly, DM-T-1 and DM-T-2 produced slightly different band patterns, especially between 25 and 37 kDa. The most intense 110 kDa band migrated at slightly higher molecular weight when glycosylated by DM-T-1 instead of DM-T-2, indicating that T-1 potentially labeled more sites of this particular protein. Digestion with the mucin-selective protease StcE completely removed this band in T-2 labeled samples (Fig. S7C), confirming labeling of mucin-type proteins that are rich in O-GalNAc glycosylation (*45*). Discrete band patterns were obtained from in-gel fluorescence experiments when GALE-KO cells were used and a functional bump-and-hole pair was present. In contrast, pulsing the same cells with GlcNAc analog **9**, a precursor of UDP-GlcNAc analog **8**, led to a diffuse background whenever mut-AGX1 was present, but independent of the GalNAcT construct used (Fig. S7D). To profile the identity of labeled cell surface glycoproteins, we synthesized clickable, acid-cleavable biotin-picolyl azide **10** as an enrichment handle of cell surface glycoproteins (Fig. 5D). Following lysis and PNGase F treatment, O-glycoproteins were enriched and digested on-bead with trypsin to profile the corresponding unmodified peptides by mass spectrometry. The myelogenous K-562 cell line expresses mucins and mucin-like proteins that are rich in O-GalNAc glycans such as CD36, CD43, CD45, Glycophorins A and C, and MUC18 (https://www.proteinatlas.org/). We found all these proteins enriched (log prob > 5 for DM and < 5 for WT samples, or at least 40 units higher for DM than controls) from lysates of cells carrying functional T-1 or T-2 bump-and-hole systems (**5**, mut-AGX1, DM-GalNAcT) over control cells lacking any component (Fig. S8). It is noteworthy that we found mainly cell surface glycoproteins with abundant O-glycosylation in these experiments, probably because DM-GalNAcTs were in constant competition for protein substrates with the corresponding endogenous WT-GalNAcTs in these cells.

The use of GalNAcT-KO SimpleCells has enabled the mapping of glycosylation sites that are introduced by several GalNAcTs including T-1 and T-2, but by design is not suited to profile the diversity of glycan structures beyond the initiating GalNAc. Our bump-and-hole approach can contribute this information if the bumped GalNAc analog is recognized and extended by downstream GTs. To probe this, we used bump-and-hole pairs to profile the glycoproteome of T-1-KO or T-2-KO HepG2 cells (*21*). Cells were transfected with GalNAcT-1 or T-2 and Dox-induced before labeling with GalNAc-1-phosphate analog **5**. As glycoproteins are abundant in the HepG2 secretome (*21*), we clicked acid-labile enrichment handle **10** onto proteins in conditioned cell culture supernatants. Following on-bead tryptic digest to release non-glycosylated peptides, glycopeptides were liberated by acidic cleavage of the enrichment handle, and analyzed by mass spectrometry. We used the presence of a 491.2238 m/z ion in higher-energy collisional dissociation (HCD) as an indicator for a bumped glycan to trigger peptide sequencing by electron-transfer dissociation (ETD, fig. 5E and fig. S9). Following an automated search for glycopeptides containing the GalNAc analog alone or as a part of longer glycans, we obtained raw glycopeptide hits which we manually validated. Seventy-seven spectra were found of DM-T-2-modified glycopeptides that corresponded to 37 peptides with 27 glycosylation sites and the presence of up to four different glycan compositions per glycopeptide, the structures of which were inferred based on biosynthetic considerations (Data S1) (*18*). In contrast, four glycopepetides were found as modified by DM-T-1. Control cells fed with DMSO or containing WT GalNAcTs and fed with GalNAc-1-phosphate analog **5** showed no hits at all, or one (WT-T-2) and two (WT-T-1) glycopeptides, respectively. Several known glycosylation sites in the SimpleCell glycoproteomics dataset by Schjoldager et al. were confirmed herein, and new sites were revealed (Data S1) (*19*). For instance, we found hitherto unannotated T-2 glycosylation sites in the proteins ITIL2, AMBP, Complement C3 and C4, GPC3 and STC2, and Laminin subunit gamma-1, among others (Fig. 5E and Data S1). T-2 is involved in lipid homeostasis and was found to glycosylate certain apolipoproteins, including ApoE at Thr307 and Ser308, in HepG2 cells (*17, 21*). Both sites were confirmed herein and, additionally, found to be elaborated up to the trisaccharide Neu5Acα-Galβ-GalNAc-Ser/Thr or tetrasaccharide Neu5Acα-Galβ-(Neu5Acα)GalNAc-Ser/Thr structures (Data S1) (*18*). Our data resolved ambiguity about O-glycosylation of apolipoprotein A-1 (ApoA1) in the peptide sequence 220-ATEHLSTLSEK-230 (*21*): previous data showed that upon T-2-KO, glycosylation of Thr221, Thr226 and Ser228 is slightly increased, but the reason for this was elusive (*21*). Our gain-of function approach revealed that DM-T-2 glycosylated Ser225 while DM-T-1 glycosylated Thr221, indicating that T-2-KO cells lose Ser225 glycosylation that is compensated by increased glycosylation at the other sites. We attribute the smaller number of annotated glycopeptides in DM-T-1 cells versus DM-T-2 to the lower expression of T-1 versus T-2 GalNAcTs at 0.5 µg/mL Dox and/or the fact that T-1 glycosylates extracellular matrix proteins which may not be sufficiently soluble in conditioned media (Fig. 3B and fig. S3A). We repeated the experiment using DM-GalNAcTs and treatment with caged GalNAc-1-phosphate analog **5** only, and found similar results (Data S2). Such detailed analysis is facilitated by the gain-of-function chemical tagging strategy that emulates native GalNAcT activity as a result of our bump-and-hole approach.

## Discussion

The cellular glycoproteome is subject to disease-related alterations. Yet, we still lack methodology to selectively visualize, modify or sequence either a certain glycan subtype or the product of a certain GT. Bump-and-hole engineering has been artfully employed to obtain insight into the protein substrates of single kinases (*4, 46*), ADP-ribosyltransferases (*5, 6*), and methyltransferases (*37, 47, 48*), among others (*9*). Only few of these systems have been established in the living cell, as cellular delivery of bumped substrates is rarely straightforward, especially when substrates are nucleotide-based. Herein, we established the first GT bump-and-hole system in the living cell to profile GalNAcTs, the largest GT family in the human genome. The fascinating biology of GalNAcTs, with multiple isoenzymes and a profound cross-talk between glycosylation and proteolysis (*34, 49*), prompted us to develop an isoenzyme-specific chemical reporter. This approach allowed us to expand our knowledge on T-1 and T-2 glycosylation sites by adding information on the nature of the mature glycans after extension by other GTs.

The era of quantitative biology demands great specificity in tools to probe cellular processes or expressed antigens. Biological tools, including antibodies, lectins, engineered binding proteins, or silenced hydrolases have been employed to profile or isolate cellular glycans (*45, 50*). While powerful probes, these proteins suffer from various drawbacks, including the dependence of binding on the structural context or the requirement to simplify glycans before performing binding studies. Additionally, all conventional strategies for glycan pull-down or visualization rely on post-glycosylation derivatization or binding. As a consequence, information on GT isoenzyme specificity is already lost or – in the context of knock-out cell lines – may have been obscured by partial redundancy. This “specificity space” is covered by our bump-and-hole approach that enables GT- and glycosite-specific labeling. As GalNAcT engineering retains both three-dimensional structure and peptide glycosylation preferences, this strategy probes glycosylation in an unbiased fashion without context dependence. Thereby, we obtained for the first time information on both site and structure of glycans initiated by a single GT isoenzyme in a single experiment.

Taken together, we now have a strategy in hand to specifically modify GT isoenzyme-dependent glycosylation sites on the living cell.

## Supporting information

Glycoproteomics data replicate 1

Glycoproteomics data replicate 2

## Acknowledgements

The authors thank Katrine T. Schjoldager and Hans H. Wandall (both University of Copenhagen, Denmark) for HepG2-T1^-/-^ and HepG2-T2^-/-^ cells and helpful discussions. We thank Lawrence Tabak (National Institutes of Health, Bethesda, MD) for full length human GalNAc-T2 in the plasmid pCMV-NTAP, Michael Bassik (Stanford University, USA) for the K-562-spCas9 cell line, Jonathan Weissman (University of California, San Francisco, USA) for lentiviral plasmids, Jon Agirre (University of York) for help on crystal structure optimization using Privateer, and Ramón Hurtado-Guerrero (University of Zaragoza, Spain) for helpful discussions. We thank David Spiciarich, Yi-Chang Liu and Christina M. Woo for help with designing experiments.

## Author contributions

B. S., M. F. D., S. P. W., S. A. M., L. L., J. J. K., M. M. and C. R. B. designed research; B. S., S. A. M., S. P. W., M. F. D., A. J. A., D. F., L. J. S. W., L. L., D. F., J. P., M. A. G. ran experiments; B. S., S. A. M., S. P. W., M. F. D., A. J. A., D. F., L. L., M. M. and C. R. B. analysed data; J. C., J. P., K. P. and J. J. K. made reagents and cell lines and contributed protocols; B. S. and C. R. B. wrote the paper with input from all authors.

## Materials & Correspondence

Correspondence and material request should be addressed to. HepG2-T1^-/-^ and HepG2-T2^-/-^ cells are subject to a materials transfer agreement with the University of Copenhagen. Plasmids used herein are subject to materials transfer agreements with addgene. Crystal structures are available in a public depository (PDB 6E7I and PDB 6NQT) and are provided with the manuscript. Mass spectrometry sequencing data will be made available in a public repository.

## Supplementary Materials

Materials and Methods

**Fig. S1.**
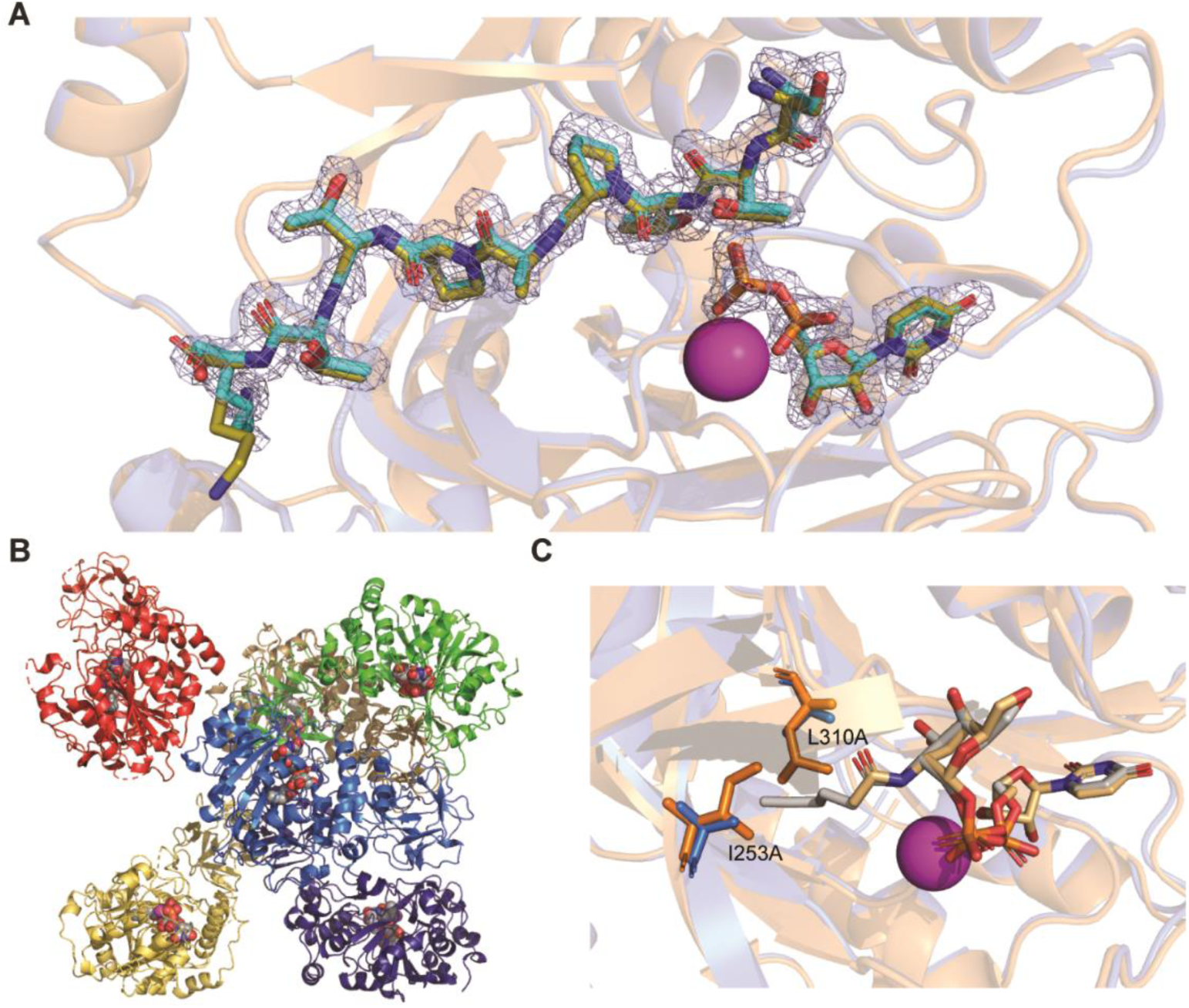
*A*, overlay of ligands bound in DM-T-2/EA2/UDP/Mn^2+^ (ligands cyan) and a published crystal structure of WT-T-2/EA2/UDP/Mn^2+^ (ligands olive, PDB 2FFU), with electron density (map rendered at 1 σ and carved at 1.6 Å) taken from the DM-T-2 co-crystal structure. Mn^2+^ ions are colored in magenta and overlay completely. *B*, crystal structure of DM-T-2/1/Mn^2+^ in the hexameric unit cell. UDP-GalNAc analog **1** is rendered in sphere representation. *C*, superposition of the active sites of WT-T-2 (PDB 4D0T, orange) with UDP-GalNAc (light brown) and DM-T-2 (blue) with UDP-GalNAc analog **1** (grey). Gatekeeper residues are rendered in stick representation. Mn^2+^ ions are magenta and overlay completely.

**Fig. S2.**
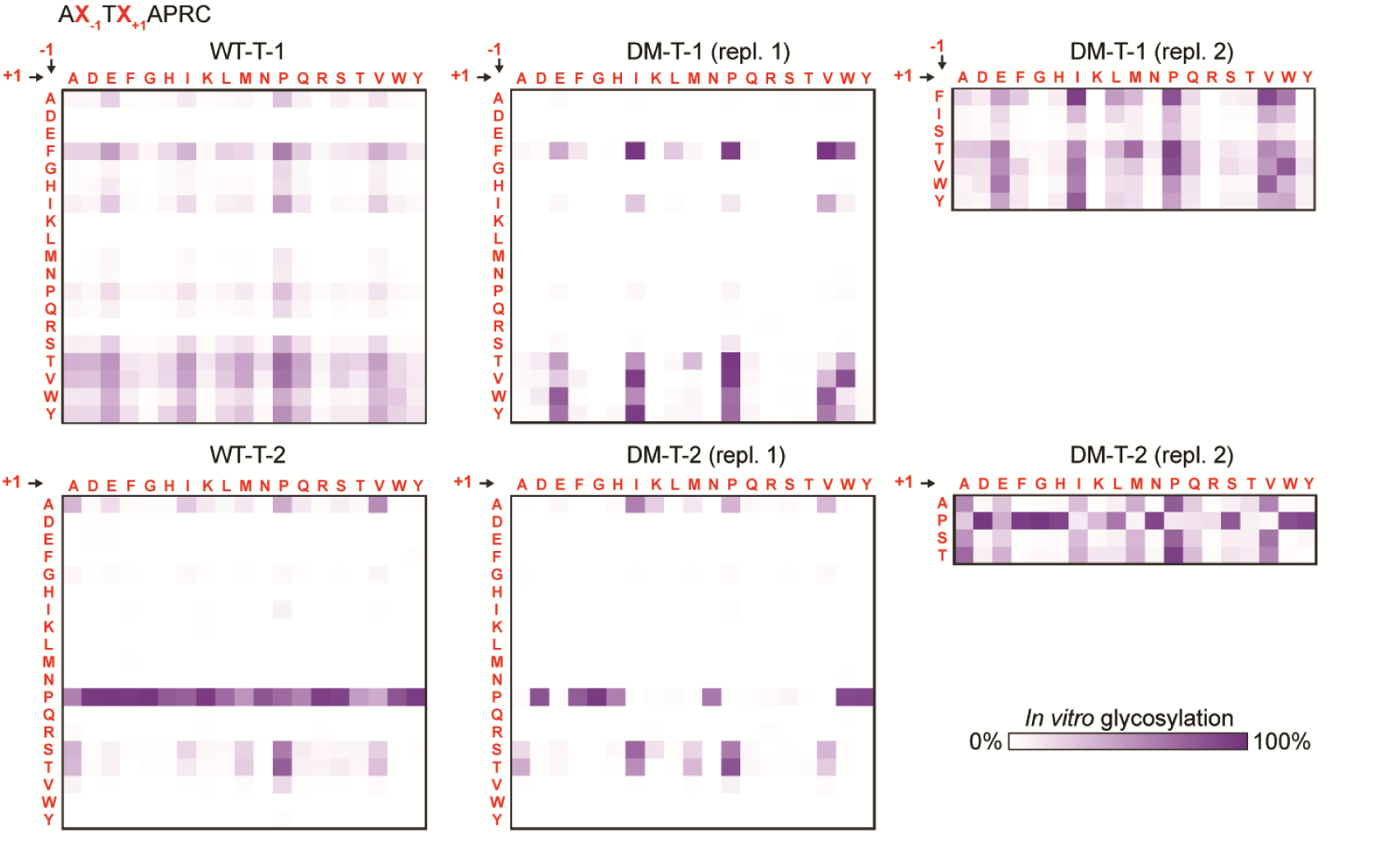
Substrate specificities of DM-T1 and DM-T-2 and comparison with WT enzymes as determined in an *in vitro* glycosylation assay with detection by SAMDI-MS. WT data corresponds with reported substrate specificities, and two replicates are shown for DM enzymes with the full library or a focused sub-library.

**Fig. S3.**
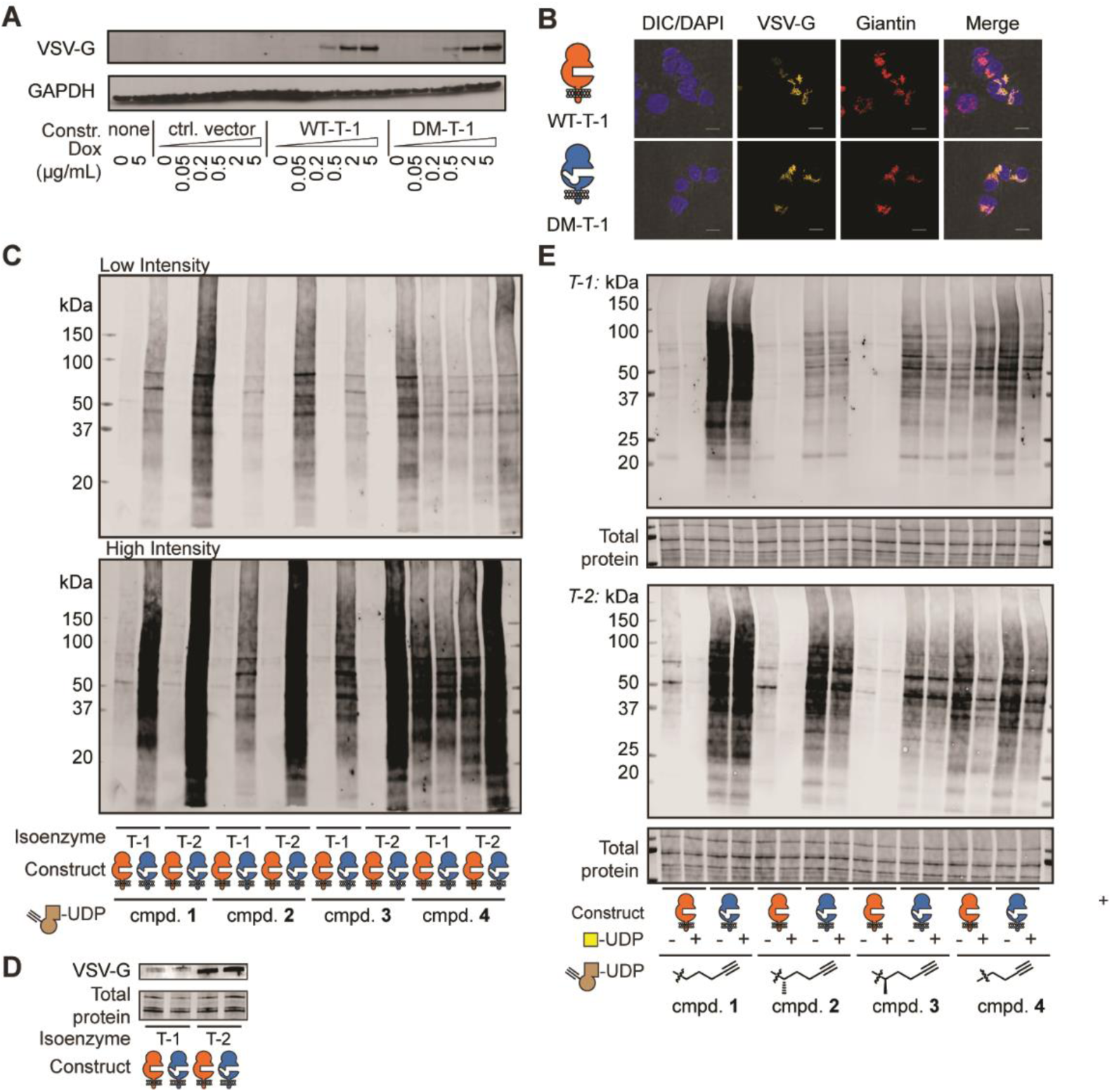
*A*, WT-T-1 and DM-T-1 are expressed by stably transfected HepG2 cells in a Dox-inducible fashion. *B*, fluorescence microscopy of HepG2 cells stably transfected with T-1 constructs, induced with 2 µg/mL Dox and subsequently stained. *C*, streptavidin blot from figure 3C depicted in two different intensities. *D*, Western blot to assess expression of GalNAcTs in lysates used in *C. E, in vitro* glycosylation was repeated with a membrane fraction from untransfected HepG2 cells, using soluble GalNAcTs to perform glycosylations (*8*). Reactions were performed with or without a two-fold excess of UDP-GalNAc over UDP-sugars **1, 2, 3** and **4**.

**Fig. S4.**
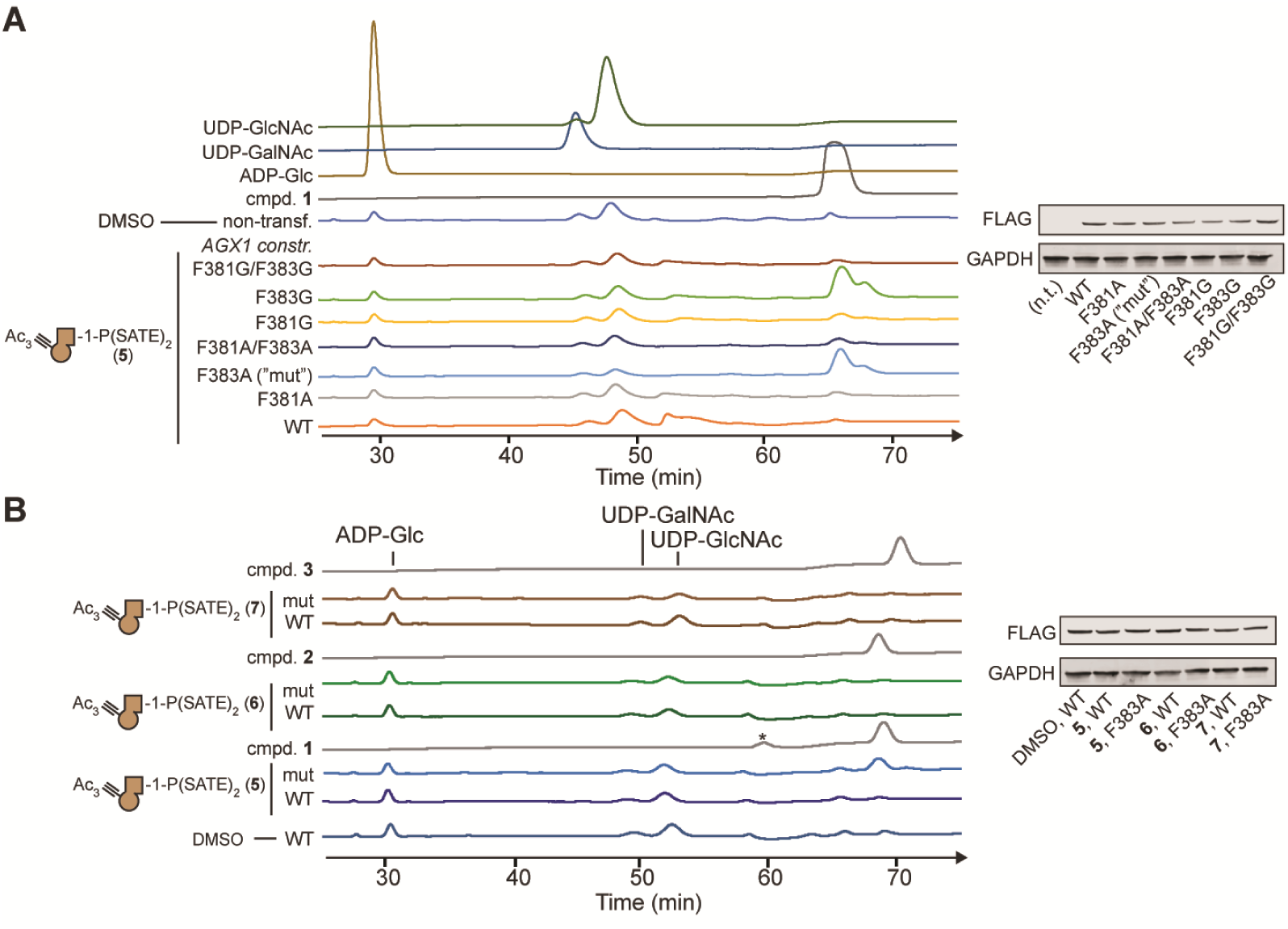
Biosynthesis of UDP-GalNAc analog **1** by mut-AGX1. *A*, HPAEC-PAD traces of extracts from HEK293T cells transiently expressing AGX1 constructs, and fed with GalNAc-1-phosphate analog **5** or DMSO. All traces are normalized to the retention time of ADP-Glc as an external standard. Data is of a single experiment. *B*, full traces of the data displayed in Fig. 4B. Asterisk indicates an artefact from solvent filling.

**Fig. S5.**
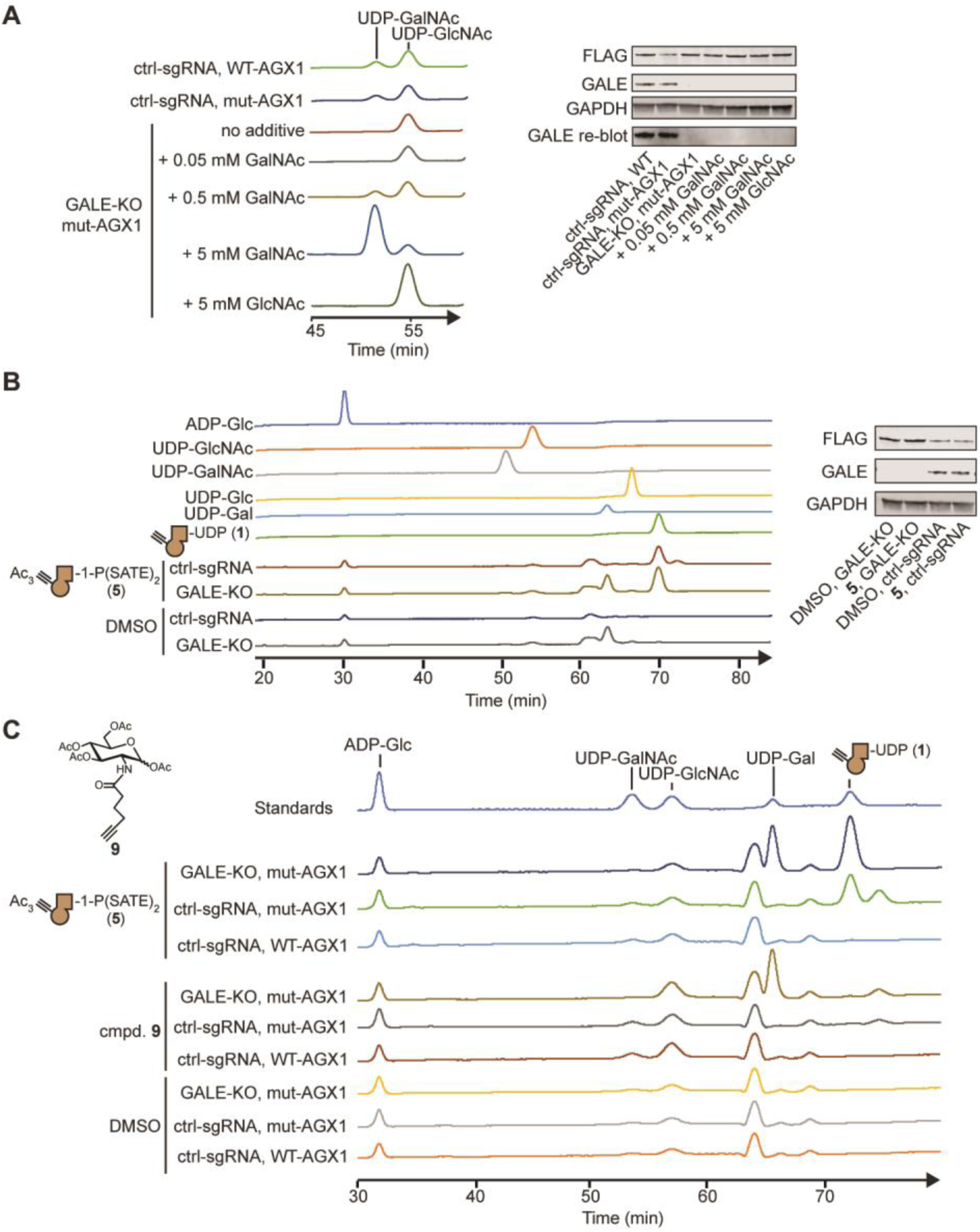
GALE-KO cells are deficient in epimerization of UDP-GalNAc and UDP-GlcNAc, as well as chemically modified analogs. *A*, HPAEC-PAD traces of GALE-KO or control sgRNA-transduced K-562 cells stably transfected with the indicated FLAG-tagged AGX1 constructs and fed with different concentrations of GalNAc or GlcNAc. Expression of AGX1 (FLAG) and GALE were analyzed by Western blot. Samples were re-blotted with a higher concentration of GALE antibody to assure absence of GALE in KO cells. Data are from one experiment. *B*, cells were fed with DMSO or compound **5**, and UDP-sugar production was measured by HPAEC-PAD. GALE-KO contain elevated levels of UDP-Gal as cells are supplemented with galactose to maintain viability and UDP-Gal cannot be epimerized to UDP-Glc. Expression levels of AGX1 (FLAG) and GALE are analyzed by Western blot. Data are of one representative out of two independent experiments. *C*, GALE-KO or control sgRNA-transduced K-562 cells stably transfected with the indicated AGX1 constructs were fed with the indicated compounds, and UDP-sugar production was measured by HPAEC-PAD. Data are from one experiment.

**Fig. S6.**
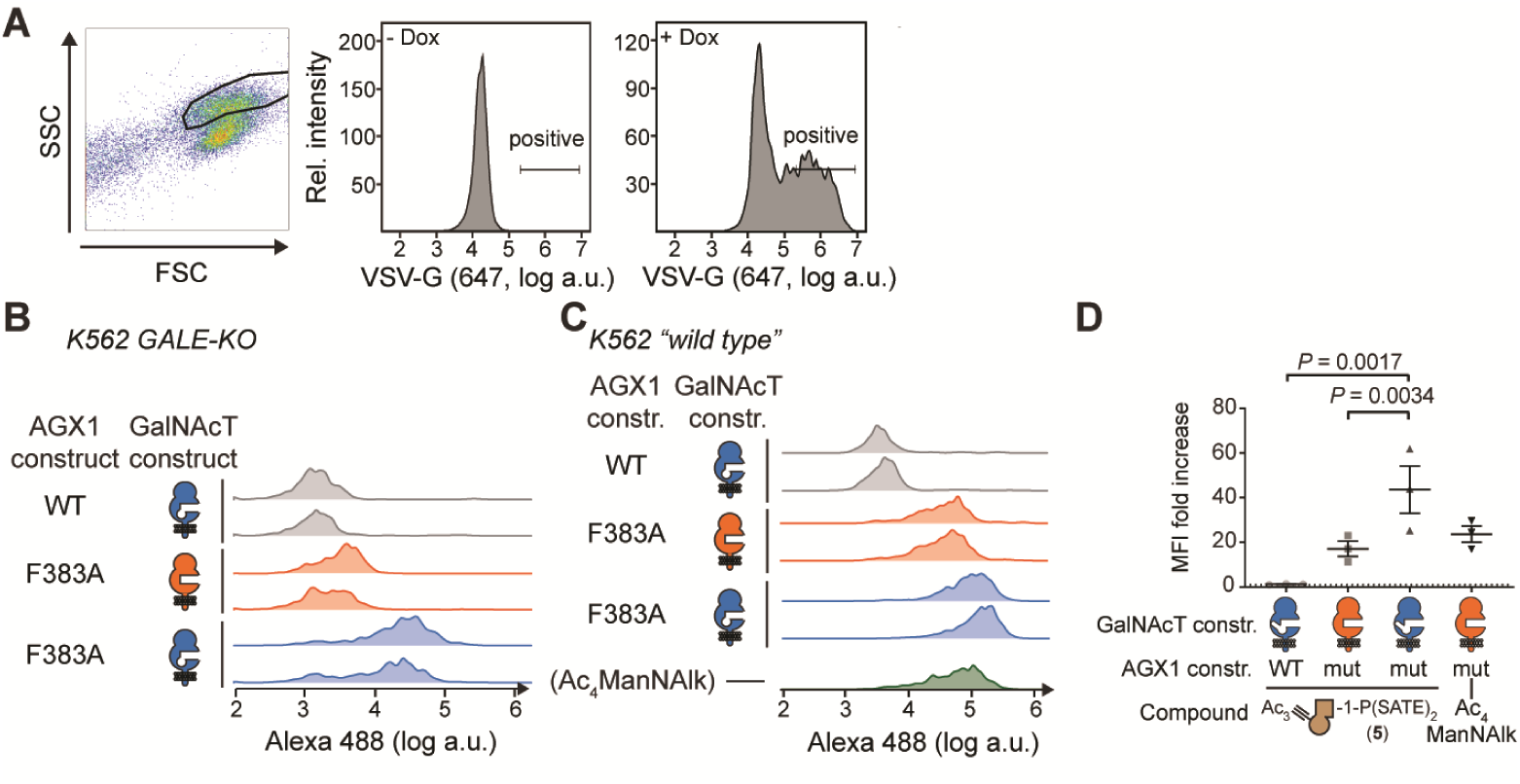
*A*, gating scheme for flow cytometry experiments in Fig. 5. *B*, primary flow cytometry data of the experiment in figure 5C. Two technical replicates are shown. *C*, primary flow cytometry data of “wild type” K-562 cells after induction with 0.5 µg/mL Dox and gating on VSV-G-positive cells. Two technical replicates are shown for cells treated with compound **5**. At least 500 gated cells were used for analysis per sample in *B* and *C. D*, statistical analysis of the experiment in *C*. Data are individual values from three independent experiments, means ± SEM of MB488 median fluorescence intensity of VSV-G positive cells. Statistical analysis was performed by two-tailed ratio paired t-test.

**Fig. S7.**
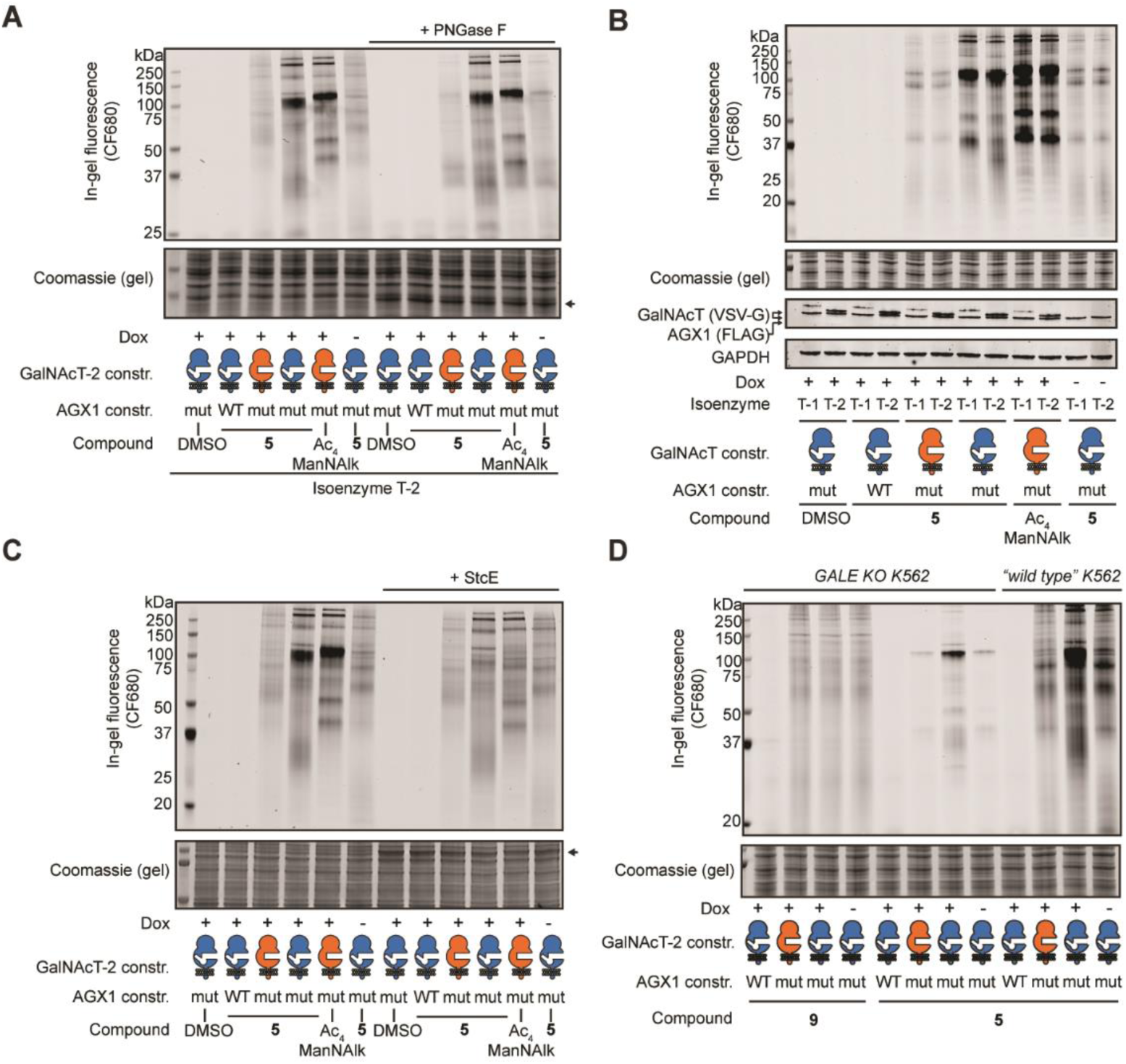
Labeling analysis of K-562 cells by in-gel fluorescence. *A*, cells expressing AGX1 and GalNAcT-2 constructs were labeled as in figure 5C and analyzed by in-gel fluorescence. Treatment of lysates with PNGase F significantly shifts certain background bands to lower molecular weight. Arrow indicates PNGase F band in Coomassie stain. *B*, full gel of the experiment depicted in figure 5C. The sialic acid precursor Ac4ManNAlk was used as a positive control, and the effect of omitting Dox was investigated. *C*, treatment of lysates prepared as in *A* with the glycoprotease StcE. Arrow indicates StcE band in Coomassie stain. *D*, dissecting GlcNAc vs. GalNAc labeling by using probes **5** and **9**. GALE-KO K-562 cells expressing AGX1 and GalNAcT-2 constructs were treated with Dox or left untreated, and fed with compounds **5** or **9**. Labeling was performed as in figure 5C. Wild type K-562 cells prepared as in *A* were used to compare labeling patterns. Data are from one representative out of three independent experiments.

**Fig. S8.**
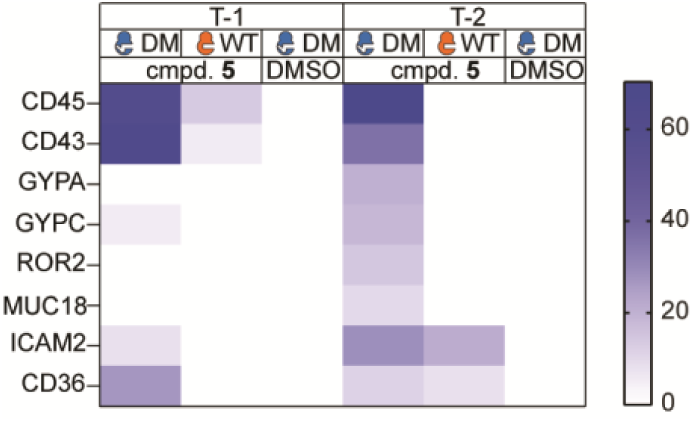
GalNAcT bump-and-hole pairs selectively label highly O-glycosylated proteins. Heat map represents log prob of detected proteins after enrichment of cell surface proteins of K-562 cells expressing mut-AGX1 and indicated GalNAcT constructs and fed with the indicated compounds.

**Fig. S9.**
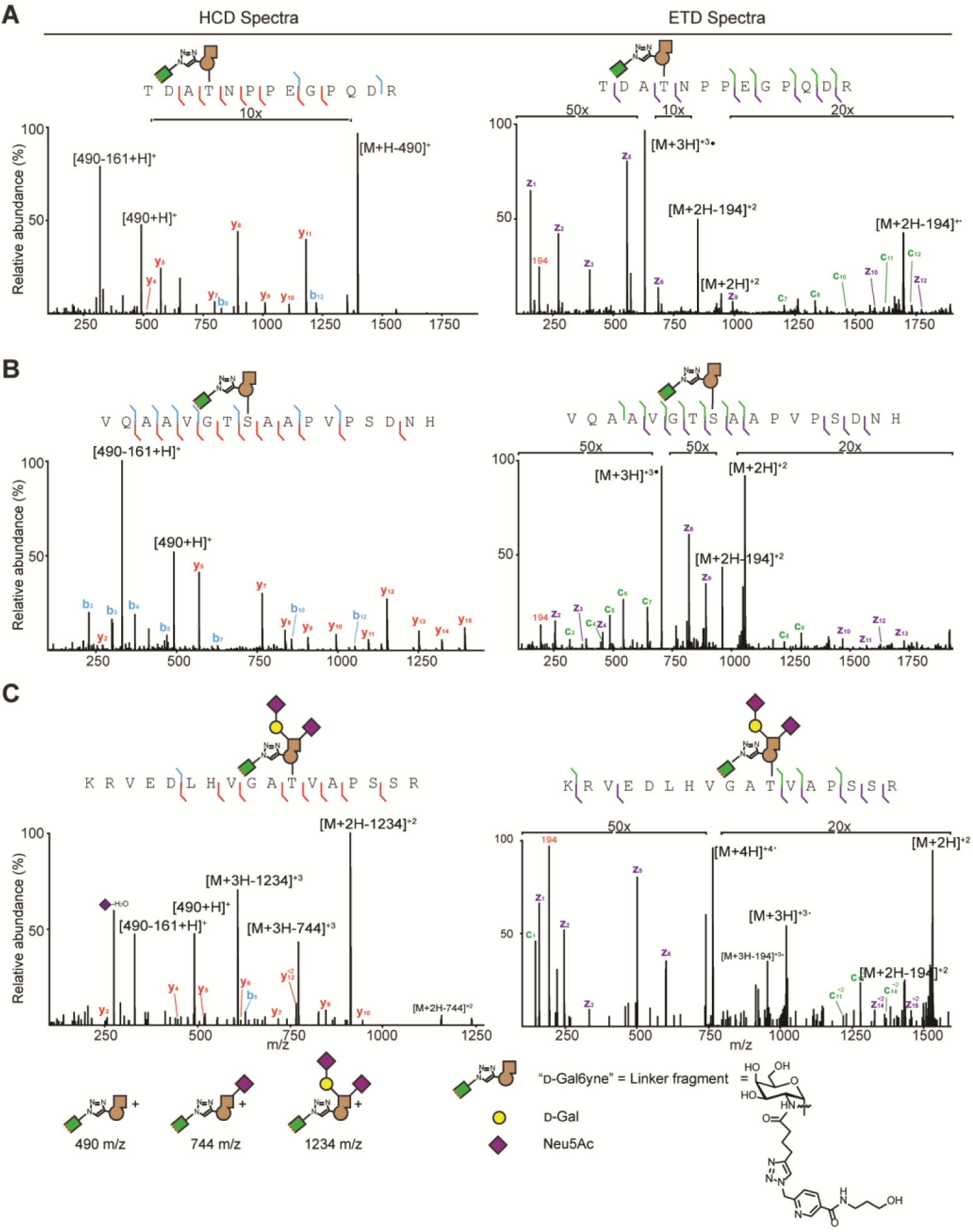
Exemplary mass spectra from glycopeptides after modification by DM-T-2. HCD (mainly glycan fragmentation) and ETD (mainly peptide fragmentation) spectra are shown, and ions are annotated. *A*, glycosylation at Thr28 of STC2 as a newly-identified T-2-specific modification (*63*). *B*, confirmation of Ser308 as a T-2-specific modification of ApoE. *C*, extension of chemically tagged GalNAc by elaborating glycosyltransferases on T39 of SERPIN5A. Legend depicts tentative structural assignments of oxonium ions. Loss of 161 m/z in HCD and 194 m/z peak in ETD likely depicts fragment masses of the triazole-based linker.

**Table S1.**
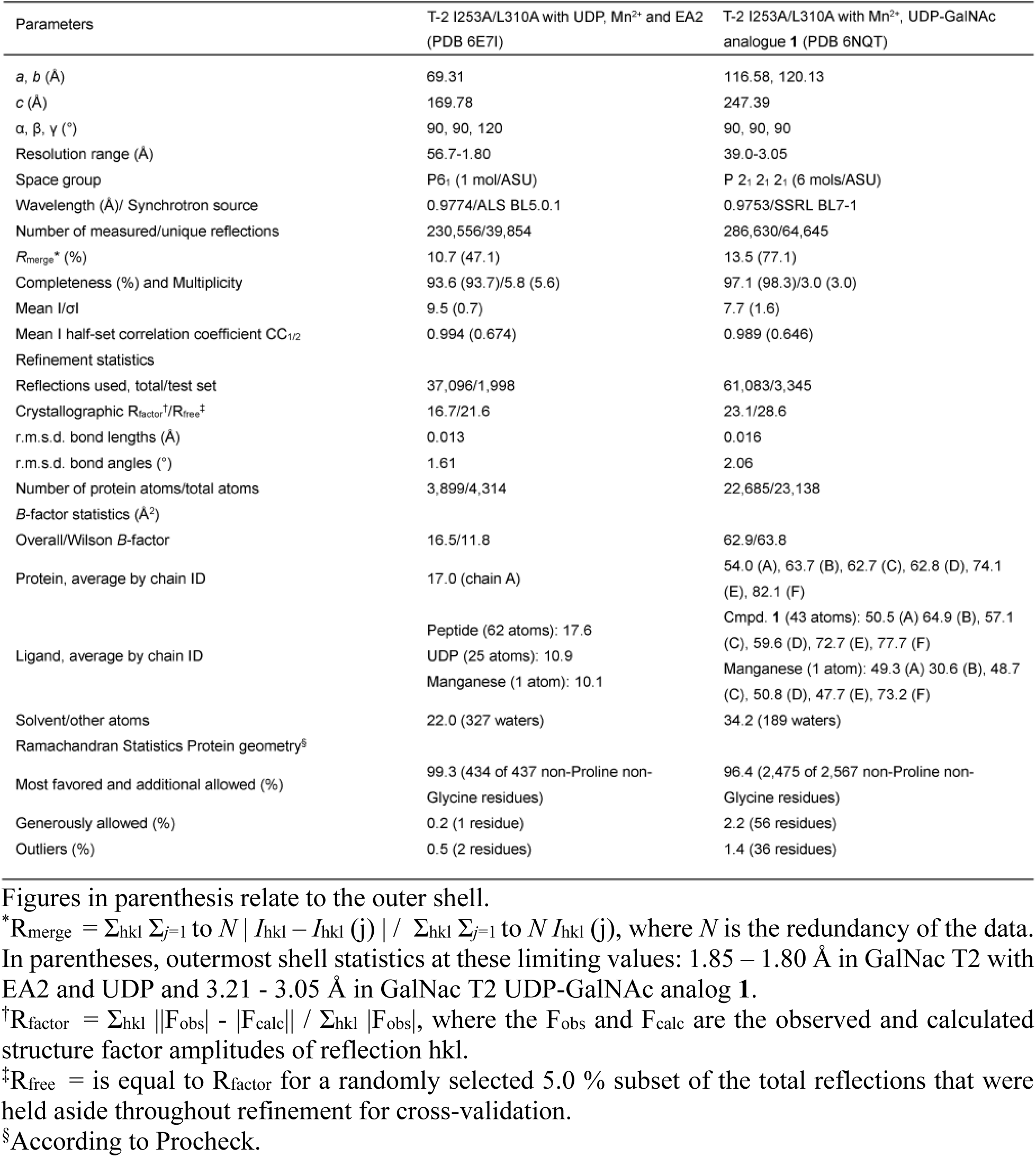
Crystallographic data statistics.

**Data S1.** Glycoproteomics data of secretome from HepG2-T1^-/-^ and HepG2-T2^-/-^ cells stably transfected with combinations of AGX1 and T-1 or T-2, respectively, and treated with compound **5** or DMSO.

**Data S2.** Glycoproteomics data of secretome from HepG2-T1^-/-^and HepG2-T2^-/-^cells stably transfected with mut-AGX1 and DM-T-1 or -T-2, respectively, and treated with compound **5**.

## References and Notes

1. A. Varki, Biological roles of glycans. Glycobiology 27, 3–49 (2017).

2. A. Varki et al., Eds, Essentials of Glycobiology, 3rd edition (Cold Spring Harbor Laboratory Press, New York, NY, 2017).

3. C. Besanceney-Webler et al., Increasing the efficacy of bioorthogonal click reactions for bioconjugation: A comparative study. Angew. Chem. Int. Ed. 50, 8051–8056 (2011).

4. P. J. Alaimo, M. A. Shogren-Knaak, K. M. Shokat, Chemical genetic approaches for the elucidation of signaling pathways. Curr. Opin. Chem. Biol. 5, 360–367 (2001).

5. I. Carter-O’Connell, H. Jin, R. K. Morgan, L. L. David, M. S. Cohen, Engineering the substrate specificity of ADP-ribosyltransferases for identifying direct protein targets. J. Am. Chem. Soc. 136, 5201–5204 (2014).

6. B. A. Gibson et al., Chemical genetic discovery of PARP targets reveals a role for PARP-1 in transcription elongation. Science. 353, 45–50 (2016).

7. K. Islam, W. Zheng, H. Yu, H. Deng, M. Luo, Expanding Cofactor Repertoire of Protein Lysine Methyltransferase. ACS Chem. Biol. 679–684 (2011).

8. J. Choi et al., Engineering orthogonal polypeptide GalNAc-Transferase and UDP-Sugar pairs. Preprint at https://chemrxiv.org/articles/Engineering_Orthogonal_Polypeptide_GalNAc-Transferase_and_UDP-Sugar_Pairs/8123486 (2019).

9. K. Islam, The Bump-and-Hole Tactic: Expanding the Scope of Chemical Genetics. Cell Chem. Biol. 25, 1171–1184 (2018).

10. E. P. Bennett et al., Control of mucin-type O-glycosylation: A classification of the polypeptide GalNAc-transferase gene family. Glycobiology 22, 736–756 (2012).

11. H. C. Hang, C. R. Bertozzi, The chemistry and biology of mucin-type O-linked glycosylation. Bioorg. Med. Chem. 13, 5021–5034 (2005).

12. A. Antonopoulos et al., Loss of effector function of human cytolytic T lymphocytes is accompanied by major alterations in N- and O-Glycosylation. 287, 11240–11251 (2012).

13. P. Radhakrishnan et al., Immature truncated O-glycophenotype of cancer directly induces oncogenic features. Proc. Natl. Acad. Sci. U. S. A. 111, E4066–75 (2014).

14. Peng, R. Q. et al., MicroRNA-214 suppresses growth and invasiveness of cervical cancer cells by targeting UDP-N-acetyl-alpha-D-galactosamine:polypeptide N-acetylgalactosaminyltransferase 7. J. Biol. Chem. 287, 14301–14309 (2012).

15. A. Gaziel-Sovran et al., MiR-30b/30d Regulation of GalNAc Transferases Enhances Invasion and Immunosuppression during Metastasis. Cancer Cell 20, 104–118 (2011).

16. M. de las Rivas, E. Lira-Navarrete, T. A. Gerken, R. Hurtado-Guerrero, Polypeptide GalNAc-Ts: from redundancy to specificity. Curr. Opin. Struct. Biol. 56, 87–96 (2019).

17. C. Steentoft et al., Mining the O-glycoproteome using zinc-finger nuclease-glycoengineered SimpleCell lines. Nat. Methods 8, 977–982 (2011).

18. K. T.-B. G. Schjoldager et al., Probing isoform-specific functions of polypeptide GalNAc-transferases using zinc finger nuclease glycoengineered SimpleCells. Proc. Natl. Acad. Sci. U. S. A. 109, 9893–8 (2012).

19. C. Steentoft et al., Precision mapping of the human O-GalNAc glycoproteome through SimpleCell technology. EMBO J. 32, 1478–1488 (2013).

20. Narimatsu, Y. et al., Exploring regulation of protein O-Glycosylation in isogenic human HEK293 cells by differential O-Glycoproteomics. Mol. Cell Proteomics https://www.mcponline.org/content/early/2019/04/30/mcp.RA118.001121 (2019).

21. K. T.-B. G. Schjoldager et al., Deconstruction of O-glycosylation-GalNAc-T isoforms direct distinct subsets of the O-glycoproteome. EMBO Rep. 16, 1713–1722 (2015).

22. J. Hintze et al., Probing the contribution of individual polypeptide GalNAc-transferase isoforms to the O-glycoproteome by inducible expression in isogenic cell lines. J. Biol. Chem. 293, 19064–19077 (2018).

23. E. M. Sletten, C. R. Bertozzi, Bioorthogonal chemistry: Fishing for selectivity in a sea of functionality. Angew. Chem. Int. Ed. 48, 6974–6998 (2009).

24. J. A. Prescher, D. H. Dube, C. R. Bertozzi, Chemical remodelling of cell surfaces in living animals. Nature 430, 873–878 (2004).

25. H. C. Hang, C. Yu, D. L. Kato, C. R. Bertozzi, A metabolic labeling approach toward proteomic analysis of mucin-type O-linked glycosylation. Proc. Natl. Acad. Sci. U. S. A. 100, 14846–51 (2003).

26. B. W. Zaro, Y.-Y. Yang, H. C. Hang, M. R. Pratt, Chemical reporters for fluorescent detection and identification of O-GlcNAc-modified proteins reveal glycosylation of the ubiquitin ligase NEDD4-1. Proc. Natl. Acad. Sci. 108, 8146–8151 (2011).

27. C. M. Woo, A. T. Iavarone, D. R. Spiciarich, K. K. Palaniappan, C. R. Bertozzi, Isotope-targeted glycoproteomics (IsoTaG): a mass-independent platform for intact N- and O-glycopeptide discovery and analysis. Nat. Methods 12, 561–567 (2015).

28. K. G. Ten Hagen, T. A. Fritz, L. A. Tabak, All in the family: The UDP-GalNAc:polypeptide N-acetylgalactosaminyltransferases. Glycobiology 13, 1–16 (2003).

29. T. A. Fritz, J. Raman, L. A. Tabak, Dynamic association between the catalytic and lectin domains of human UDP-GalNAc:polypeptide α-N-acetylgalactosaminyltransferase-2. J. Biol. Chem. 281, 8613–8619 (2006).

30. E. Lira-Navarrete et al., Substrate-guided front-face reaction revealed by combined structural snapshots and metadynamics for the polypeptide N-acetylgalactosaminyltransferase 2. Angew. Chem. Int. Ed. 53, 8206–8210 (2014).

31. E. Lira-Navarrete et al., Dynamic interplay between catalytic and lectin domains of GalNAc-transferases modulates protein O-glycosylation. Nat. Commun. 6, 6937–6948 (2015).

32. W. Kightlinger, et al., Design of glycosylation sites by rapid synthesis and analysis of glycosyltransferases. Nat. Chem. Biol. 14, 627–635 (2018).

33. T. A. Gerken, et al., Emerging paradigms for the initiation of mucin-type protein O-glycosylation by the polypeptide GalNAc transferase family of glycosyltransferases. J. Biol. Chem. 286, 14493–14507 (2011).

34. L. Revoredo, et al., Mucin-type O-glycosylation is controlled by short- and long-range glycopeptide substrate recognition that varies among members of the polypeptide GalNAc transferase family. Glycobiology 26, 360–76 (2015).

35. S. Ji, et al., A molecular switch orchestrates enzyme specificity and secretory granule morphology. Nat. Commun. 9, 3508 (2018).

36. E. Kowarz, D. Löscher, R. Marschalek, Optimized sleeping beauty transposons rapidly generate stable transgenic cell lines. Biotechnol. J. 10, 647–653 (2015).

37. R. Wang et al., Profiling genome-wide chromatin methylation with engineered posttranslation apparatus within living cells. J. Am. Chem. Soc. 135, 1048–1056 (2013).

38. M. Boyce et al., Metabolic cross-talk allows labeling of O-linked β-N-acetylglucosamine-modified proteins via the N-acetylgalactosamine salvage pathway. Proc. Natl. Acad. Sci. USA 108, 3141–3146 (2011).

39. S. Pouilly, V. Bourgeaux, F. Piller, V. Piller, Evaluation of analogs of GalNAc as substrates for enzymes of the Mammalian GalNAc salvage pathway. ACS Chem. Biol. 7, 753–760 (2012).

40. S.-H. Yu et al., Metabolic labeling enables selective photocrosslinking of O-GlcNAc-modified proteins to their binding partners. Proc. Natl. Acad. Sci. 109, 4834–4839 (2012).

41. J. M. Schulz et al., Determinants of function and substrate specificity in human UDP-galactose 4′-epimerase. J. Biol. Chem. 279, 32796–32803 (2004).

42. C. Uttamapinant et al., Fast, cell-compatible click chemistry with copper-chelating azides for biomolecular labeling. Angew. Chem. Int. Ed. 51, 5852–5856 (2012).

43. W. Qin et al., Artificial cysteine S-glycosylation induced by per-O-acetylated unnatural monosaccharides during metabolic glycan labeling. Angew. Chem. Int. Ed. 57, 1817–1820 (2018).

44. P. V. Chang et al., Metabolic labeling of sialic acids in living animals with alkynyl sugars. Angew. Chem. Int. Ed. 48, 4030–4033 (2009).

45. S. A. Malaker et al., The mucin-selective protease StcE enables molecular and functional analysis of human cancer-associated mucins. Proc. Natl. Acad. Sci. USA doi:10.1073/pnas.1813020116 (2019).

46. A. C. Bishop et al., A chemical switch for inhibitor-sensitive alleles of any protein kinase. Nature 407, 395–401 (2000).

47. K. Islam et al., Defining efficient enzyme-cofactor pairs for bioorthogonal profiling of protein methylation. Proc. Natl. Acad. Sci. USA 110, 16778–16783 (2013).

48. J. Li, H. Wei, M. M. Zhou, Structure-guided design of a methyl donor cofactor that controls a viral histone H3 lysine 27 methyltransferase activity. J. Med. Chem. 54, 7734–7738 (2011).

49. S. L. King et al., TAILS N-terminomics and proteomics reveal complex regulation of proteolytic cleavage by O-glycosylation. J. Biol. Chem. 293, 7629–7644 (2018).

50. M. Chen et al., An engineered high affinity Fbs1 carbohydrate binding protein for selective capture of N-glycans and N-glycopeptides. Nat. Commun. 8, 1–15 (2017).

51. W. Kabsch, XDS. Acta Cryst. D 66, 125–132 (2010).

52. P. Evans, Scaling and assessment of data quality. Acta Cryst. D 62, 72–82 (2006).

53. M. D. Winn, et al., Overview of the CCP4 suite and current developments. Acta Cryst. D 67, 235–242 (2011).

54. L. C. Storoni, et al., Accelerating structural biology with Phaser crystallographic software. Acta Cryst. 5711, 1373–1382 (2001).

55. T. A. Gerken, J. Raman, T. A. Fritz, O. Jamison, Identification of common and unique peptide substrate preferences for the UDP-GalNAc:polypeptide α-N-acetylgalactosaminyltransferases T1 and T2 derived from oriented random peptide substrates. J. Biol. Chem. 281, 32403–32416 (2006).

56. P. Emsley, B. Lohkamp, W. G. Scott, K. Cowtan, Features and development of Coot. Acta Cryst. D 66, 486–501 (2010).

57. G. N. Murshudov, A. A. Vagin, E. J. Dodson, Refinement of macromolecular structures by the maximum-likelihood method. Acta Cryst. D 53, 240–255 (1997).

58. V. B. Chen et al., MolProbity: All-atom structure validation for macromolecular crystallography. Acta Cryst. D 66, 12–21 (2010).

59. J. Agirre et al., Privateer: Software for the conformational validation of carbohydrate structures. Nat. Struct. and Mol. Biol. 22, 833–834 (2015).

60. J. M. Termini et al., HEK293T cell lines defective for O-linked glycosylation. PLoS One 12, 1–19 (2017).

61. L. A. Gilbert et al., CRISPR-mediated modular RNA-guided regulation of transcription in eukaryotes. Cell 154, 442–451 (2012).

62. L. Mátés et al., Molecular evolution of a novel hyperactive Sleeping Beauty transposase enables robust stable gene transfer in vertebrates. Nat. Genet. 41, 753–761 (2009).

63. C. W. Luo, M. D. Pisarska, A. J. W. Hsueh, Identification of a stanniocalcin paralog, stanniocalcin-2, in fish and the paracrine actions of stanniocalcin-2 in the mammalian ovary. Endocrinology 146, 469–476 (2005).

64. N. S. Berrow et al., A versatile ligation-independent cloning method suitable for high-throughput expression screening applications. Nucleic Acids Res. 35, pdoi:10.1093/nar/gkm047 (2007).

65. T. Dull et al., A third-generation lentivirus vector with a conditional packaging system. J. Virol. 72, 8463–8471 (1998).

