## Supplementary material for "Chemical precision glyco-mutagenesis by glycosyltransferase engineering in living cells": Materials and Methods

**This PDF file includes:**

Materials and Methods
Figs. S1 to S9
Table S1
Captions for Data S1 and S2

**Other Supplementary Materials for this manuscript include the following:**

Data S1 and S2

### Materials and Methods

The following nomenclature of GalNAcT and AGX1 genes will be used herein: WT-T-1, WT-T-2: wild type constructs; DM-T-1: I238A/L295A; DM-T-2: I253A/L310A; mut-AGX1: F383A.
All cell lines were mycoplasma-free, as confirmed using the MycoAlert Kit (Lonza, Basel, Switzerland).

Cloning of full length GalNAcT-1 and T-2

Primers and gBlocks were from Integrated DNA Technologies (IDT, Coralville, USA) and Elim Biopharm (Hayward, USA). Restriction enzymes were from New England Biolabs except for PasI (Thermo Fisher, Waltham, USA). PfuUltra II (Agilent, Santa Clara, USA) and Advantage Genomic LA polymerase (Takara, Kusatsu, Japan) were used for T-1 and T-2, respectively. T4 DNA ligase (New England Biolabs, Ipswitch, USA) and LONG DNA ligase (Takara) were used for ligation of T-1 and T-2, respectively. Q5 high-fidelity DNA polymerase (New England Biolabs) was used for overlap extension PCR. All plasmids were sequenced before use at Elim Biopharmaceuticals, Inc. (Hayward, USA).
The plasmid pSBtet-GH was a gift from Eric Kowarz (Addgene plasmid #60498; http://n2t.net/addgene:60498 ; RRID:Addgene_60498) (*36*).
Full length human GalNAcT-2 (EBI accession number LC043140.1) in pCMV-NTAP was a kind gift from Lawrence Tabak (National Institutes of Health, Bethesda, MD) (*8*). T-2 was first sub-cloned with a C-terminal FLAG tag into a version of pFLAG-CMV-2 (Sigma-Aldrich) in which the N-terminal FLAG tag had been removed, using a two-step PCR elongation procedure with the primers CTTGCGGCCGCGATGCGGCGGCGCTCGCGGATGC (fwd for both steps), GTCATCGTCTTTGTAGTCCTGCTGCAGGTTGAGCGTGAACTTCCACTGC (rev step 1) and GATGAATTCCTACTTGTCGTCATCGTCTTTGTAGTCCTGCTGCAGGTTGAGCG (rev step 2), and a NotI/EcoRI restriction strategy. Site-directed mutagenesis (I253A/L310A) was performed at this step according to a published procedure (*8*). T-2 constructs were cloned into pFLAG-CMV-5.1 (Sigma Aldrich) using the primers GCAGAGCTCGTTTAGTGAACCGTCAGAATTGATCTAC (fwd) and TCTCTCGGATCCCTGCTGCAGGTTGAGGGTGAACTTCC (rev), and a NotI/BamHI strategy. These constructs were used as templates to clone T-2 constructs into pSBtet-GH (*36*) with a C-terminal VSV-G tag, using the primers AAAGGCCTCTGAGGCCACCATGCGGCGGCGCGCTCG (fwd) and TTTGGCCTGACAGGCCCTACTTACCCAGGCGGTTCATTTCGATATCAGTGTACTGCTGCAGGTTGAGCGGTG (rev) with an SfiI cloning strategy.
Full length GalNAcT-1 (NCBI Genbank® accession number NM_020474) was used as a gBlock (IDT) with overhangs and NotI/BamHI restriction sites: AAATTTGCGGCCGCAGTGCCATGAAAATATTCGGATCCGGGCCC and amplified with the primers AAATTTGCGGCCGCAGTGCC (fwd) and GGGCCCGGATCCGAATATTTCTGG (rev) prior to cloning into pFLAG-CMV-5.1 using a NotI/BamHI strategy. Site-directed mutagenesis (I238A/L295A) was performed at this step according to a published procedure (*8*). These constructs were used as templates to clone T-1 constructs into pSBtet-GH with a C-terminal VSV-G tag, using the primers AAAGGCCTCTGAGGCCACCATGAGAAAATTTGCATACTGCAAG (fwd) and TTTGGCCTGACAGGCCCTACTTACCCAGGCGGTTCATTTCGATATCAGTGTAGAATATTTCTGGCAGGGTGACGTTTC (rev) with an SfiI cloning strategy.
Cloning, expression and crystallization of DM-T-2

The plasmid pOPING was a gift from Ray Owens (Addgene plasmid # 26046 ; http://n2t.net/addgene:26046; RRID:Addgene_26046) (*64*). A secretion construct of DM-T-2 (aa 75-571), chosen according to literature precedent of crystallization of the WT enzyme (*30*), with a $His_6$ tag in pOPING was used for protein expression and crystallisation. The coding sequence in pFLAG-Myc-CMV-19 (*8*) was cloned into KpnI/PmeI-digested pOPING using the primers

GCGTAGCTGAAACCGGCAAAGTACGGTGGCCAGACTTTAACCAG (fwd) and GTGATGGTGATGTTTCTGCTGCAGGTTGAGCGTGAA (rev), with In-Fusion HD Cloning Kit (Takara). Freestyle™ 293-F cells were propagated in FreeStyle™ 293 Expression Medium (Thermo Fisher) with 10% (v/v) fetal bovine serum (FBS, Thermo Fisher) at 37 °C and 8% $CO_2$ with orbital rotation at 135 rpm. Cells were transfected with pOPING DM-T-2 using 293fectin (Thermo Fisher) according to the manufacturer's specifications, with $5x10^7$ to $1x10^8$ cells and 30 μg plasmid DNA. After 24 h, cells were harvested (5 min, 500 g, room temperature) and medium was renewed (30 mL). Cells were harvested and conditioned supernatant collected after one, three, and five days, and medium was renewed each time. Conditioned supernatant was passed through a freshly-packed column containing 2 mL HisPur NiNTA resin slurry (Thermo Fisher) pre-conditioned with water and Wash Buffer (20 mM imidazole, 50 mM Tris-HCl, 125 mM NaCl, pH 7.5). The resin was treated with Wash Buffer (2x15 mL), and subsequently with Elution Buffer (12.5 mL, 200 mM imidazole, 50 mM Tris-HCl, 125 mM NaCl, pH 7.5, cOmplete protease inhibitors (Roche, Basel, Switzerland)). Eluted protein was dialyzed using Amicon Ultra-15 centrifuge filters (10 kDa MWCO, Millipore) against Crystallization Buffer (25 mM Tris-HCl, 0.5 mM EDTA, 1 mM TCEP, pH 8.0). Protein was stable for one week at 4 °C. For long-term storage at -80 °C, 80% (v/v) aq. glycerol was added to a final concentration of 25% (v/v). Protein was re-buffered to Crystallization Buffer immediately prior to use. Typically, 1.2 mg protein was purified from one 30 mL sample of conditioned supernatant.

Crystals were grown using sitting drop vapour diffusion at room temperature. Drops were set-up using a Douglas Oryx8 Nanodrop Dispensing Robot (Douglas Instruments Ltd, Berkshire, United Kingdom).

Crystals of DM-T-2 complexed with UDP, $Mn^{2+}$ and EA2 peptide were obtained through co-crystallization using 0.15 μL of 5 mM $MnCl_2$, 5 mM UDP, 6 mM EA2, and 10 mg/mL protein in Crystallization Buffer and 0.15 μL precipitant solution (11-32% (v/v) PEG8000, 0.1 mM HEPES, pH 6.5-8.5) against 80 μL of precipitant solution. Crystals were grown for 1 week before being cryoprotected in 16% ethylene glycol and frozen in liquid nitrogen before diffraction.

Crystals of DM-T-2 complexed with **1** and $Mn^{2+}$ were obtained through soaking of EA2 and **1** into co-crystals of DM-T2 with UDP and $Mn^{2+}$. EA2 peptide electron density was too weak to properly discern in crystal structures. Initial drops were created using 0.15 μL of 5 mM $MnCl_2$, 5 mM UDP, 10 mg/mL T2 DM protein in 25 mM Tris-HCl (pH 8.0), 0.5 mM EDTA, and 1 mM TCEP and 0.15 μL precipitant solution (11-23% PEG8000, 0.1 mM HEPES, pH 7.5-8.5) against 80 μL of precipitant solution. Crystals were grown for 18 days before soaking and diffraction. Crystals were soaked in a solution with EA2 by adding 0.45 μL of a 50 mM EA2 solution in water to the original drop for 30 minutes to 2 hours. Crystals were harvested and soaking was performed with a solution of 20 mM **1** with 16% ethylene glycol, 11-23% PEG8000, 0.1 mM HEPES, 5 mM $MnCl_2$, pH 7.5-8.5 for 1-2 hours.

Single crystal X-ray diffraction of DM-T-2/EA2/UDP/$Mn^{2+}$ (PDB 6E7I) was performed at 95K using ALS Beamline 5.0.1 (Lawrence Berkeley National Labs, Berkeley, USA) and single wavelength of 0.97741 Å, with a Dectris Pilatus3 S 6M Detector (Dectris Ltd., Baden-Daettwil, Switzerland).

Single crystal X-ray diffraction of DM-T-2/**1**/$Mn^{2+}$ (PDB 6NQT) was performed at 100K using Stanford Synchrotron Radiation Lightsource Beamline 7.1 (SLAC National Accelerator Laboratory, Menlo Park, CA, USA) and single wavelength of 0.9753 Å, with an ADSC Quantum 315r CCD Detector (Quantum Detectors Ltd, Harwell Oxford, United Kingdom).

Data was processed using XDS (*51*), and scaled with SCALA (*52*) and other programs implemented with the CCP4 software (*53*). Crystal structures were determined using molecular replacement with Phaser (*54*), using published structures (PDB 2FFU for 67EI and PDB 4D0T for 6NQT) as the templates (*29, 30*). The initial model was improved through multiple cycles of manual modeling in Coot (*56*), and refinement using REFMAC5 (*57*). The final structural model was validated using MOLPROBITY (*58*). UDP-GalNAc analog conformation was validated using Privateer (*59*). Images were prepared using Pymol 2.0.0 (Schrödinger LLC, New York, USA). Electron density was rendered at 1 σ and carved at 1.6 Å. Structure statistics are given in Table S1.

### Screening peptide substrate selectivities of GalNAcTs using SAMDI-MS

SAMDI-MS screenings were performed similarly to previously reported (*32*). Briefly, 50 mM solutions of 361 peptides of the general formula $AX_{-1}TX_{+1}APRC$, where $X_{-1}$ and $X_{+1}$ are 19 natural amino acids except for Cys, were screened individually with 25 nM purified WT-T-1 or 50 nM purified WT-T-2 and 0.25 mM UDP-GalNAc in SAMDI buffer (50 mM Tris-HCl pH 7.4, 50 mM NaCl, 5 mM $MnCl_2$), at 37 °C for 3 h (*8*). For DM-GalNAcTs, the same peptide array was screened with 30 nM purified DM-T-1 and 0.25 mM compound **1** in buffer 1 at 37 °C for 21 h, or with 175 nM DM-T-2 and 0.25 mM compound **1** in buffer 1 at 37 °C for 3 h (*8*). After the reaction of each 10 µL volume, 5 mM EDTA were added for quenching and 2.5 µL TCEP reducing gel (Thermo Fisher) were added for reduction incubation for 1 h at 37 °C. Reduced mixture (2 µL) was added to the gold islands of a 384 SAMDI plate, prepared as described before (*32*). The SAMDI plate was incubated at room temperature for 30 min, and washed with water, ethanol and water, nitrogen blow dried, and treated with 10 mg/mL 2',4',6'-trihydroxyacetophenone monohydrate (Sigma-Aldrich) matrix in acetonitrile. The SAMDI plates were analyzed with a Sciex MALDI-TOF/TOF 5800 (Applied Biosystems, Foster City, USA), equipped with the software Explorer 4.1.0 to analyse data. %Intensity of glycosylated peptides related to the sum of unreacted and glycosylated peptide was recorded. Preferred $X_{-1}$ amino acids, combined with all $X_{+1}$ amino acids, were chosen to repeat the experiment for DM-GalNAcTs.

### Cloning and site-directed mutagenesis of AGX1

$AGX1^{F383G}$ in pIRES-puro3 was used as a template to generate AGX1 constructs (*40*). Site-directed mutagenesis was performed using the following primers (positions 381 and 383 underlined): CAAACCCAATGGAATAAAGATGGAAAAAGGTGTCTTTGACATCTTCCAG (fwd) and CAAAGACACCTTTTTCCATCTTTATTCCATTGGGTTTGTCTGGCTTAATTAAC (rev) for F381G; CAAACCCAATGGAATAAAGATGGAAAAAGGTGTCGGTGACATCTTCCAG (fwd) and CACCGACACCTTTTTCCATCTTTATTCCATTGGGTTTGTCTGGCTTAATTAAC (rev) for F381G/F383G. A C-terminal FLAG-tag was then introduced into these constructs using a site-directed mutagenesis strategy with the primers (FLAG sequence underlined) GACTACAAAGACGATGACGACAAGTGAGCGGCCGCATAGATAACTGATCC (fwd) and CTTGTCGTCATCGTCTTTGTAGTCAATACCATTTTTCACCAGCTCATGAACTCCATTCTC (rev). These constructs were then used to produce other AGX1 constructs. Primer pairs (positions 381 and 383 underlined) were: GGAAAAAGCTGTCTTTGACATCTTCCAGTTTGC (fwd) and GATGTCAAAGACAGCTTTTTCCATCTTTATTCCATTGGG (rev) for F381A; GAATAAAGATGGAAAAATTTGTCGCTGACATCTTCCAGTTTGC (fwd) and GATGTCAGCGACAAATTTTTCCATCTTTATTCCATTGGGTTTG (rev) for F383A; GATGGAAAAAGCTGTCGCTGACATCTTCCAGTTTGC (fwd) and GATGTCAGCGACAGCTTTTTCCATCTTTATTCCATTGGG (rev) for F381A/F383A; CCCAATGGAATAAAGATGGAAAAATTTGTCTTTGACATCTTCC (fwd) and GACAAATTTTTCCATCTTTATTCCATTGGGTTTGTCTGGC (rev) for WT.

### Cloning of AGX1 into pSBtet plasmids

WT-AGX1 or mut-AGX1 were cloned into the backbone of pSBtet-GH containing WT-T-1, DM-T-1, WT-T-2, or DM-T-2 by replacing the eGFP coding sequence. C-terminally FLAG-tagged AGX1 sequences were amplified from the corresponding pIRES-puro3 constructs (see above) using the primers ATGAACATTAATGACCTCAAACTCACGTTGTCC (fwd) and CTTGTCATCGTCTTTGTAGTCAATACCATTTTTC (rev), purified by gel extraction, and subjected to an overlap extension PCR with equimolar amounts (10 fmol each) of the gBlocks (IDT) GGCCCGCCTTCCCTGGGGAATCTCTGCGCACGCGCAGAACGCTTCGACCAATGAAAACACAGGAAGCCGTCCGCGCAACCGCGTTGCGTCACTTCTGCCGCCCCTGTTTCAAGGTATATAGCCGTAGACGGAACTTCGCCTTTCTCTCGGCCTTAGCGCCATTTTTTTGGGTGAGTGTTTTTTGGTTCCTGCGTTGGGATTCCGTGTACAATCCATAGACATCTGACCTCGGCACTTAGCATCATCACAGCAAACTAACTGTAGCCTTTCTCTCTTTCCCTGTAGAAACCTCTGCACCTGAGGCCACCATGAACATTAATGACCTCAAACTCACGTTGTCCAAAGCTGGGC (N-gBlock) and CATGAGCTGGTGAAAAATGGTATTGACTACAAAGACGATGACGACAAGGGCAGTGGAGCTACTAACTTCAGCCTGCTGAAGCAGGCTGGTGACGTCGAGGAGAATCCTGGCCCCATGTCTAGACTGGACAAGAGCAAAGTCATAAACGGC (C-gBlock) using 10 mM dNTPs and Q5 polymerase. Briefly, a 50 µL reaction was run for 15 cycles using the following protocol: 95 °C (2 min), 95 °C (30 s), 64 °C (30 s), 72 °C (30 s – return to step 2), 72 °C (5 min), then treated with the primers (500 nmol) GGCCCGCCTTCCCTGG (fwd) and GCTCTTGTCCAGTCTAGACATGGGGC (rev) and continued for another 18 cycles with an annealing temperature of 55 °C. The resulting PCR product was used to replace the eGFP cloning sequence of pSBtet-GH containing T-1 or T-2 using an XcmI/PasI strategy to digest the vector and In-Fusion HD Cloning Kit to insert AGX1 sequences. AGX1-containing pSBtet vectors are named pSBtet-WT-hAGX1 or pSBtet-mut-hAGX1.

pSBtet-based vectors herein are pSBtet-GH (eGFP, Luciferase), pSBtet-WT-hAGX (WT-AGX1, Luciferase), pSBtet-mut-AGX1 (mut-AGX1, Luciferase), pSBtet-GH-WT-T-1 (eGFP, WT-T-1), pSBtet-GH-DM-T-1 (eGFP, DM-T-1), pSBtet-GH-WT-T-2 (eGFP, WT-T-2), pSBtet-GH-DM-T-2 (eGFP, DM-T-2), pSBtet-mut-hAGX1-WT-T-1 (mut-AGX1, WT-T-1), pSBtet-WT-hAGX1-DM-T-1 (WT-AGX1, DM-T-1), pSBtet-DM-hAGX1-DM-T-1 (DM-AGX1, DM-T-1), pSBtet-mut-hAGX1-WT-T-2 (mut-AGX1, WT-T-2), pSBtet-WT-hAGX1-DM-T-2 (WT-AGX1, DM-T-2), pSBtet-DM-hAGX1-DM-T-2 (DM-AGX1, DM-T-2).

### Generation of K-562 GALE$^{-/-}$ cells

K-562 cells with stable expression of *Streptococcus pyogenes* Cas9 (K-562-spCas9) were a gift from Michael Bassik (Stanford University, USA) and propagated in RPMI (Thermo Fisher) with 10% (v/v) FBS, penicillin (100 U/mL) and streptomycin (100 µg/mL). Expression of spCas9 was confirmed by Western blot. Single-guide (sg) RNAs targeting human *GALE* were chosen according to Termini et al. with the sequences CCGGGATTACATCCATGTCG (sgRNA_GALE1) and TCAGCTCCTGGACCCGCCGC (sgRNA_GALE2) (*60*). Specificity and minimal off-target effects of sgRNAs were further confirmed *in silico* (www. synthego.com). SgRNAs were cloned into the vector pU6-sgGAL4-4 (*61*), a gift from Jonathan Weissman (University of California, San Francisco), using a BstXI-BlpI restriction strategy and annealed oligodeoxynucleotides with the following overhangs as inserts: TTG-[forward sgRNA]-GTTTAAGAGC and TTAGCTCTTAAAC-[reverse complementary sgRNA]-CAACAAG.

HEK293T cells were seeded at 1.2 million cells per well of a 6-well plate. After 24 h, cells were transfected with 1.5 µg sgRNA-coding vector and a third generation lentiviral packaging mix (0.1 µg GAG/POL, 0.1 µg REV, and 0.2 µg VSV-G (*65*), using TransIT-293 (Mirus Bio LLC, Madison, USA). After 16 h, supernatant was aspirated and replaced with fresh growth medium. After another 30 h, supernatant was collected, and filtered by 0.45 µm syringe filtration. Virus-containing supernatant was stored at -80 °C.

For lentiviral infection, 200,000 cells were seeded in 1 mL medium in a 24-well plate. After 24 h, 300 µL virus-containing supernatant and 8 µL polybrene (Sigma) were added. After another 24 h, cells were harvested and treated with fresh growth medium. After another 24 h, cells were harvested and treated with growth medium containing 1.25 µg/mL puromycin. Selection was carried out for 96 h, and cells were expanded in puromycin-free growth medium. The knockout cells were fed 100 uM of GalNAc for 2 days before FACS sorting. After 48 h, the cells were stained with 5 µg/mL biotinylated *Vicia villosa* lectin (Vector Labs Burlingame, USA) and 4 µg/mL of AlexaFluor488 conjugated streptavidin (Invitrogen) on ice for 30 min. The cells were washed twice with PBS + 0.5% BSA and viable cells were sorted based on AlexaFluor488 fluorescence intensity and low SYTOX Red (Thermo) staining according to a published procedure (*15*). GALE-KO cells were maintained in standard growth medium supplemented with 20 µM galactose and 200 µM GalNAc. GALE was not detectable by Western blot using mouse anti-GALE sc-390407 (Santa Cruz Biotechnology, Dallas, USA) over at least 20 passages. UDP-GalNAc was absent in these cells unless GalNAc was supplemented (Fig. S6A). Both sgRNAs had the same effect, and only sgRNA_GALE2 was used for further experiments.

Cell transfection

Tet-system approved FBS (Takara) was used to propagate all cell lines transfected with pSBtet-GH-based plasmids. pCMV(CAT)T7-SB100 was a gift from Zsuzsanna Izsvak (Addgene plasmid # 34879 ; http://n2t.net/addgene:34879 ; RRID:Addgene_34879) (*62*).

HEK293T (ATCC CRL-3216) were grown in DMEM (Thermo Fisher) with 10% (v/v) FBS (Thermo Fisher), penicillin (100 U/mL) and streptomycin (100 µg/mL, GE Healthcare, Chicago, USA). Cells were transfected with pIRES-puro3 plasmids containing AGX1 constructs using TransIT-293 (Mirus Bio LLC, Madison, USA) according to the manufacturer's instructions and 37.5 µg DNA per 15 cm dish or 15 µg DNA per 10 cm dish. After 24 h, medium was aspirated, and cells were either treated with fresh growth medium and compounds for analysis of nucleotide-sugar biosynthesis (see below), or growth medium containing 5 µg/mL puromycin (Sigma-Aldrich) for selection of stable cells for two weeks.

K-562 cells were a gift from Jonathan Weissman (University of California, San Francisco). Cells were grown in RPMI with 10% (v/v) FBS, penicillin (100 U/mL) and streptomycin (100 µg/mL). Cells were transfected with pSBtet-based plasmids using Lipofectamine LTX (Thermo Fisher) according to the manufacturer's specifications, with 200000 cells in 1 mL growth medium, 1.25 µg pSBtet and 62.5 ng pCMV(CAT)T7-SB100 plasmid DNA. After 24 h, cells were harvested and selected in growth medium containing 150 µg/mL hygromycin B (Thermo Fisher) for 7-10 days to obtain stable cells.

HepG2 cells (ATCC HB-8065), HepG2-T1$^{-/-}$ and HepG2-T2$^{-/-}$ cells (a gift from Katrine T. Schjoldager and Hans H. Wandall, University of Copenhagen, Denmark) were propagated in low-glucose DMEM (Caisson Labs, Smithfield, USA) with 10% (v/v) FBS, penicillin (100 U/mL) and streptomycin (100 µg/mL). Cells were transfected with Lipofectamine 3000 (Thermo Fisher) according to the manufacturer's instructions, e.g. using 2.5 µg pSBtet and 125 ng pCMV(CAT)T7-SB100 plasmid DNA per well of a 6-well plate. After 24 h, medium was aspirated, and cells were treated with fresh growth medium containing 600 µg/mL (HepG2) or 500 µg/mL (HepG2-T1$^{-/-}$ and HepG2-T2$^{-/-}$) hygromycin B for two weeks to obtain stable cells. Following selection, cells were propagated in 200 µg/mL hygromycin B in growth medium.

GalNAcT expression analysis and membrane lysate labeling

HepG2 cells stably transfected with pSBtet-GH, pSBtet-GH-T1$^{WT}$, pSBtet-GH-T1$^{DM}$, pSBtet-T2$^{WT}$, or pSBtet-T2$^{DM}$ were seeded into 6-well plates at 40% confluency. After 24 h, the supernatant was aspirated and changed to fresh growth medium without hygromycin B, containing the indicated concentration of doxycycline (AppliChem, Maryland Heights, USA). After 24 h, the treatment was repeated in fresh growth medium. After another 24 h, cells were washed with PBS (1 mL), treated with ice-cold PBS containing 1 mM EDTA (1 mL) and incubated for 10 min at 4 °C. Cells were transferred to a 1.5 mL reaction tube, harvested (600 g, 5 min, 4 °C), and resuspended in ice-cold 50 mM Tris-HCl pH 8, 5 mM EDTA, 150 mM NaCl, 1% (v/v) NP-40, 0.5% (v/v) sodium deoxycholate, 0.1% (w/v) SDS (0.2 mL) containing cOmplete protease inhibitors (Roche, Basel, Switzerland). Cells were briefly vortexed, incubated for 30 min at 4 °C with agitation, and centrifuged (18000 g, 30 min, 4 °C). The supernatant was transferred to a new tube, and the protein concentration was measured by BCA (Thermo Fisher). SDS-PAGE and Western blot were performed using 15 µg protein per lane, and blots were decorated with mouse anti-VSV-G-tag P5D4 (abcam, Cambridge, UK) and rabbit anti-GAPDH-HRP conjugate ab185059 (abcam).

Membrane protein glycosylation was performed on a membrane fraction from untransfected or stable HepG2 cells. Cells were seeded in a 15 cm dish, grown to 50% confluency and treated with doxycycline (5 µg/mL final concentration). After 24 h, doxycycline treatment was repeated. After another 24 h, cells were washed with PBS (10 mL) and scraped in PBS (5 mL). Cells were harvested by centrifugation (300 g, 5 min, 4 °C), washed once with ice-cold PBS (2 mL) and frozen at -80 °C. Cells were thawed on ice and fractionated using the Subcellular Fractionation Kit for Cultured Cells (Thermo Fisher) according to the manufacturer's instructions, with 0.4 mL of both Cytoplasmic Extraction Buffer and Membrane Extraction Buffer. Protein concentration was measured by BCA (Thermo Fisher), and fractions were frozen after addition of 80% (v/v) glycerol (10% final concentration).

*In vitro* glycosylation reactions were performed using 10 µg membrane protein of stably transfected HepG2 cell lines in 20 µL reaction volume containing 62.5 mM Tris-HCl pH 7.4, 150 mM NaCl, 10 mM $MnCl_2$, 250 µM UDP-GalNAc analog and 500 µM UDP-GalNAc at 37 °C for 12 h. Reactions were heat-inactivated at 95 °C for 20 s and cooled to 4 °C. Then, alkyne-containing reaction mixtures were sequentially treated with equal volumes (1.25 µL each) of 2 mM biotin-$PEG_4$-azide (Thermo Fisher), 2 mM BTTAA (Click Chemistry Tools, Scottsdale, USA), 20 mM $CuSO_4$ and 100 mM sodium ascorbate (final concentrations 100 µM biotin probe, 100 µM BTTAA, 1 mM $CuSO_4$ and 5 mM sodium ascorbate). Click reactions were carried out at room temperature for 2 h and quenched by addition of 50 mM EDTA. Reaction mixtures were then subjected to SDS-PAGE and blotted on nitrocellulose membranes. The total protein amount was assessed using the REVERT protein staining kit (LI-COR Biosciences, Lincoln, USA), and biotinylation was detected using IRDye 800CW Streptavidin (LI-COR Biosciences) according to the manufacturer's instructions.

*In vitro* glycosylations were replicated using a membrane protein fraction from untransfected HepG2 cells that was depleted for internal GalNAcT activity by three heat (90 °C)-cool (4 °C) cycles of 30 s each. Soluble WT or DM GalNAcTs (*8*) were added at a final concentration of 20 nM (T-1) or 10 nM (T-2), along with bumped UDP-GalNAc analog (250 µM) and UDP-GalNAc (500 µM). Glycosylation, click reaction and Streptavidin detection were performed as described above.

Fluorescence microscopy

Fluorescence microscopy was performed with stably transfected HepG2 cells. Cells were Dox-induced (T-1-transfected cells: 2 µg/mL; T-2-transfected cells: 0.2 µg/mL) for 24 h, then trypsinated and transferred in growth medium to a 24 well plate containing circular 12 mm coverslips (Electron Microscopy Services, Hatfield, USA) pre-treated with human fibronectin (5 µg/mL in PBS, Merck & Co., Kenilworth, USA) for 20 min and washed with PBS. Cells were treated with Dox again and incubated for another 24 h. Supernatant was aspirated, cells were washed in PBS with 100 mg/L $Ca^{2+}$ and 100 mg/mL $Mg^{2+}$ (DPBS), and fixed with 4% (v/v) paraformaldehyde (Sigma) in PBS for 10 min in the dark. Cells were washed with 50 mM ammonium chloride in DPBS and DPBS with 2% (w/v) BSA (Incubation Buffer). Cells were then permeabilized with 0.5% (v/v) Tween-20 in incubation buffer for 10 min in the dark, and subsequently treated with antibody solutions in the same buffer: mouse anti-VSV-G-tag, (1:200), and rabbit anti-Giantin (1:500, 9B6, abcam). Cells were incubated for 1 h, washed twice with Incubation Buffer with 0.1% Tween-20, and incubated with secondary antibodies in Incubation Buffer containing 0.5% (v/v) Tween-20: Donkey anti-mouse Alexa Fluor 568 (1:200, abcam) and donkey anti-rabbit Alexa Fluor 647 (1:200, abcam). Cells were incubated for 1 h in the dark, washed twice with Incubation Buffer with 0.1% (v/v) tween-20, and mounted on glass slides using ProLong Diamond Antifade mounting solution (Thermo Fisher). Cells were imaged using a Nikon A1R+ Resonant Scanning Confocal Microscope.

Analysis of nucleotide-sugar biosynthesis by High Performance Anion Exchange Chromatography

Cells expressing AGX1-FLAG (transient or stably transfected HEK293T in 20 mL growth medium in a 15 cm dish or 5 million stable K-562 in 4 mL growth medium) were treated with caged GalNAc-1-phosphate analogs (100 µM final concentration from 10 mM or 100 mM stock solutions in DMSO; all samples of an experiment contained equal amounts of DMSO) or DMSO vehicle. After 7 h, K-562 cells were harvested (500 g, 5 min, 4 °C) and the supernatant was aspirated. HEK293T cells were washed once on the plate with cold PBS (8 mL), scraped in cold 1 mM EDTA in PBS (8 mL), transferred to a conical tube and harvested (300 g, 5 min, 4 °C). Cell pellets were resuspended in PBS (1 mL), 0.8 mL of that suspension was transferred to O-ring tubes (1.5 mL, Thermo Fisher) and harvested. Zirconia/silica beads (0.1 mm, BioSpec, Bertlesville, USA) were added at a similar volume as the cell pellet, followed by 1:1 acetonitrile/water (1 mL). Cells were lysed with a bead beater (FastPrep-24, MP Biomedicals, Santa Ana, USA) for 30 s at 6 m/s, and the mixture was left at 4 °C for 10 min. Samples were centrifuged (14000 g, 10 min, 4 °C), and the resulting supernatant was transferred to a new 1.5 mL tube. The solvent was removed by speed vac. The residue was resuspended in a solution of 15 µM ADP-α-D-glucose (Sigma) in LCMS-grade water (Thermo Fisher, 0.2-0.4 mL). The solution was membrane-filtered (30 min, 14000 g) using a 3 kDa cut-off filter (Centricon, Merck) and analyzed by high performance anion exchange chromatography. The residual cell suspension in PBS (0.2 mL) was harvested, and the pellet was resuspended in M-PER lysis buffer (Thermo Fisher) containing cOmplete protease inhibitor (0.2 mL). The solution was incubated for 10 min at room temperature and centrifuged (14000 g, 10 min, 4 °C). The protein concentration of the supernatant was measured by BCA, and samples were used for analysis of protein expression.

High performance anion exchange chromatography was performed on an ICS-5000 with a quaternary pump, a CarboPac PA1 4x250 mm column with a corresponding 4x50 mm guard column and pulsed amperometric detection (Thermo Fisher). The gradient used was: A = 1 mM NaOH in degassed water; C = 1 mM NaOH, 1M NaOAc in degassed water; 0 min (A: 95; C: 5); 20 min (A: 60; C: 40); 60 min (A: 60; C: 40); 63 min (A: 50; C: 50); 83 min (A: 50; C: 50); 87 min (A: 0; C: 100); 95 min (A: 0; C: 100); 97 min (A: 95; C: 5); 105 (A: 95; C: 5). Mixtures of commercial or synthetic standards (100-400 µM) were used.

Cell surface labeling, flow cytometry and in-gel fluorescence

K-562 cells stably transfected with pSBtet-hAGX1$^{WT}$-T1$^{DM}$, pSBtet-hAGX1$^{3A}$-T1$^{WT}$, pSBtet-hAGX1$^{3A}$-T1$^{DM}$ pSBtet-hAGX1$^{WT}$-T2$^{DM}$, pSBtet-hAGX1$^{3A}$-T2$^{WT}$, pSBtet-hAGX1$^{3A}$-T2$^{DM}$ were seeded into 6-well plates at a density of 500,000 cells/2 mL in growth medium without hygromycin and treated with 0.5 µg/mL doxycycline or left untreated. Cells were treated with doxycycline again after 24 h. After another 24 h, cells were counted, harvested and seeded at a density of 50,000 cells/0.2 mL (flow cytometry) or 200,000 cells/0.6 mL (in-gel fluorescence) fresh growth medium. Cells were treated with doxycycline again, and either caged GalNAc-1-phosphate analog **5**, $Ac_4$ManNAlk (50 µM final concentration each) or DMSO vehicle. Cells were grown for another 20 h, harvested in a V-shaped 96 well plate and washed twice with 2% FBS in PBS (Cell Buffer, 0.2 mL).

For in-gel fluorescence, cells were resuspended in Cell Buffer (35 µL), treated with a solution of 50 µM $CuSO_4$, 300 µM BTTAA (Click Chemistry Tools, Scottsdale, USA), 2.5 mM sodium ascorbate, 2.5 mM aminoguanidinium chloride and 50 µM CF680 picolyl azide in cell buffer (35 µL), and incubated for 7 min at room temperature on an orbital shaker. The reaction was quenched by addition of 3 mM bathocuproinedisulfonic acid in PBS (35 µL). Cells were harvested, washed twice with Cell Buffer and once with PBS, and resuspended in ice-cold Lysis Buffer (50 mM Tris-HCl pH 8, 150 mM NaCl, 1% (v/v) Triton X-100, 0.5% (v/v) sodium deoxycholate, 0.1% (w/v) SDS, 1 mM $MgCl_2$, and 100 mU/µL benzonase (Merck) containing cOmplete protease inhibitors, 0.1 mL). Cells were lysed for 20 min at 4 °C on an orbital shaker and centrifuged (1500 g, 20 min, 4 °C). The supernatant was transferred to a new plate and protein concentration was measured by BCA. For enzymatic digest, equal amounts of protein (typically 15 µg)

were diluted to 40 µL with Lysis Buffer (PNGase F digest) or PBS (StcE digest), treated with PNGase F (2 U, Promega, Madison, USA) or StcE (50 nM) and incubated for 12 h at 37 °C (*45*). The reaction was quenched by heating to 95 °C for 10 s with subsequent cooling at 4 °C. Loading buffer (a 1:1:1:0.5 (v/v/v/v) mixture of 1 M Tris-HCl pH 7.0, 10% (w/v) SDS, 80% (v/v) glycerol and 1 M DTT) was added, samples were heated at 95 °C for one minute, run on a 10% Criterion™ gel (Bio-Rad, Hercules, USA) for SDS-PAGE, and imaged on an Odyssey CLx (LI-COR Biosciences, Lincoln, USA). Total protein content was assessed by Coomassie staining using Acquastain (Bulldog Bio, Portsmouth, USA). Another aliquot of each sample was used for protein expression control by Western blot, using antibodies against VSV-G-tag, GAPDH (ab128915, abcam) or FLAG tag (mouse anti-FLAG M2, Sigma Aldrich).
For flow cytometry, cells were resuspended in cell buffer (50 µL) and treated with a solution of 50 µM $CuSO_4$, 300 µM BTTAA, 2.5 mM sodium ascorbate, 2.5 mM aminoguanidinium chloride and 50 µM MB488 picolyl azide (Click Chemistry Tools) in Cell Buffer (50 µL), and incubated for 5 min at room temperature on an orbital shaker. The reaction was quenched by addition of 3 mM bathocuproinedisulfonic acid in PBS (50 µL). Cells were harvested, washed twice with cell buffer and once with PBS, Cells were then fixed with 0.5% (v/v) paraformaldehyde in PBS (100 µL) for 10 min at room temperature in the dark, harvested and washed with cell buffer once. Cells were permeabilized with 0.5% (v/v) Tween-20 in cell buffer (100 µL) for 10 min at room temperature. Cells were harvested, treated with mouse anti-VSV-G-tag (1:200) in 0.5% (v/v) Tween-20 in cell buffer (50 µL) for 30 min at room temperature, and washed with 0.1% (v/v) Tween-20 in cell buffer. Cells were treated with anti-mouse IgG 647 (1:200, Jackson ImmunoResearch, Cambridgeshire, UK) in 0.5% (v/v) Tween-20 in cell buffer (50 µL) for 30 min at room temperature, harvested, washed with 0.1% (v/v) Tween-20 in Cell Buffer and Cell Buffer without detergent. Flow cytometry was performed on an Accuri C6 flow cytometer (Becton Dickinson, Franklin Lakes, USA).

<u>Proteomics</u>

K-562 cells stably transfected with pSBtet-hAGX1$^{3A}$-T1$^{WT}$, pSBtet-hAGX1$^{3A}$-T1$^{DM}$, pSBtet-hAGX1$^{3A}$-T2$^{WT}$, or pSBtet-hAGX1$^{3A}$-T2$^{DM}$ were seeded into a T75 flask at 5,000,000 cells/15 mL in growth medium without hygromycin, and treated with 0.5 µg/mL doxycycline. Cells were treated with 10 mL growth medium and 0.5 µg/mL doxycycline after 24 h. After another 24 h, cells were counted, harvested and seeded at a density of 10,000,000 cells/25 mL in growth medium without hygromycin. Cells were treated with doxycycline again, and either caged GalNAc-1-phosphate analog **5** (50 µM) or DMSO vehicle. Cells were grown for another 20 h, harvested and washed with PBS (5 mL). Cell pellets were stored at -80 °C until use.
Cell pellets were resuspended in 500 µL Lysis Buffer (PBS with 1% (v/v) RapiGest and 1x Calbiochem Protease Inhibitors, Set III) and homogenized using a finger tip sonicator (output 1.0, 6x10 s strokes with 10 s breaks) on ice. Five to six mg protein per sample were treated with PNGase F (Promega, 5 µL of a 1:10 (v/v) dilution in PBS) and incubated for 4 h at 37 °C. The sample was diluted with PBS to a protein content of 2 mg/mL. Beads (UltraLink PLUS Streptavidin, 150 µL slurry = 75 µL settled resin) were washed with PBS twice (harvest 30 s at 1000 g in table top) in low-bind tubes (Eppendorf) and added to the lysate. Samples were incubated for 16 h at room temperature under rotation. The beads were harvested and the supernatant was discarded. The beads were washed sequentially with 0.1% RapiGest in PBS (3x), 6 M urea in PBS (3x), and PBS (2x), and resuspended in PBS (200 µL). Beads were treated with 10 µL 100 mM DTT (Thermo Fisher) in PBS (10 µL), and incubated at r.t. while shaking (950 rpm) for 30 min. Then, 500 mM iodoacetamide (Sigma Aldrich) in PBS (4 µL) was added and samples were shaken for another 30 min in the dark. Beads were harvested and washed with PBS and 50 mM ammonium bicarbonate in LC/MS-grade water (3x, ABC buffer). Beads were resuspended in ABC buffer (200 µL), treated with RapiGest to a final concentration of 0.05% (v/v), and trypsin (1.5 µg in 3 µL ABC buffer). Samples were shaken at 37 °C for 2-3 h, and another 1.5 µg trypsin was added. The reactions were shaken overnight at 37 °C. The beads were

harvested, washed with ABC (200 µL) and with LC-MS grade water (3x200 µL) Supernatants and washes were combined and centrifuged (18000 g, 5 min, room temperature). The supernatants were transferred to new tubes and concentrated by Speedvac. Samples were desalted by C18 Zip tips, using 0.1% aq. formic acid (FA) as washing solution and stepwise elution with 50% MeCN/water with 0.1% FA (v/v) and 100% MeCN with 0.1% (v/v) FA.

Samples were analyzed by online nanoflow LC-MS/MS using an Orbitrap Fusion Tribrid mass spectrometer (Thermo Fisher) coupled to a Dionex Ultimate 3000 HPLC (Thermo Fisher). A portion of the sample (6.5 µL out of 8 µL for glycopeptide fractions and 2 µL out of 10 µL for peptide fractions) was loaded via autosampler isocratically onto a C18 nano pre-column using 0.1% formic acid in water ("Solvent A"). For pre-concentration and desalting, the column was washed with 2% ACN and 0.1% formic acid in water ("loading pump solvent"). Subsequently, the C18 nano pre-column was switched in line with the C18 nano separation column (75 µm x 250 mm EASYSpray (Thermo Fisher) containing 2 µm C18 beads) for gradient elution. The column was held at 40 °C using a column heater in the EASY-Spray ionization source (Thermo Fisher). The samples were eluted at a constant flow rate of 0.3 µL/min using a 90 minute gradient and a 140 minute instrument method. The gradient profile was as follows (min:% solvent B, 2% formic acid in acetonitrile) 0:3, 3:3, 93:35, 103:42, 104:95, 109:95, 110:3, 140:3. The instrument method used an MS1 resolution of 60,000 at FWHM 400 m/z, an AGC target of 3e5, and a mass range from 300 to 1,500 m/z. Dynamic exclusion was enabled with a repeat count of 3, repeat duration of 10 s, exclusion duration of 10 s. Only charge states 2-6 were selected for fragmentation. MS2s were generated at top speed for 3 s. HCD was performed on all selected precursor masses with the following parameters: isolation window of 2 m/z, 28-30% collision energy, orbitrap (resolution of 30,000) detection, and an AGC target of 1e4 ions. ETD was performed if (a) the precursor mass was between 300-1000 m/z and (b) 2 of 10 glyco and/or fingerprint ions (126.055, 138.055, 144.07, 168.065, 186.076, 204.086, 274.092, 292.103; 491.2241, 330.1554) were present at +/- 0.1 m/z and greater than 5% relative intensity. ETD parameters were as follows: calibrated charge-dependent ETD times, 2e5 reagent target, precursor AGC target 1e4.

Data evaluation was performed with Byonic™ (Protein Metrics, Cupertino, USA). For protein IDs, Uniprot human proteome (downloaded June 26, 2016) was used as a reference database. Search parameters included semi-specific cleavage specificity at the C-terminal site of R and K, with two missed cleavages allowed. Mass tolerance was set at 10 ppm for MS1s, 0.1 Da for HCD MS2s, and 0.35 Da for ETD MS2s. Methionine oxidation (common 2), asparagine deamidation (common 2), and N-terminal acetylation (rare 1) were set as variable modifications with a total common max of 3, rare max of 1. Cysteine carbamidomethylation was set as a fixed modification. Peptide hits were filtered using a 1% FDR. Additionally, a cut-off value of Log Prob = 5 was set for protein hits.

For glycopeptide analysis, search parameters included semi-specific cleavage specificity at the C-terminal site of R and K, with two missed cleavages allowed. Mass tolerance was set at 10 ppm for MS1s, 0.1 Da for HCD MS2s, and 0.35 Da for ETD MS2s. Methionine oxidation (common 2), asparagine deamidation (common 2), and N-terminal acetylation (rare 1) were set as variable modifications with a total common max of 2, rare max of 1. O-glycans were also set as variable modifications (common 2), using a custom database, whereby HexNAc, HexNAc-NeuAc, HexNAc-Hex, HexNAc-Hex-NeuAc, and HexNAc-Hex-NeuAc2 were searched with an additional 287.1371 m/z to account for the chemical modification. HCD was used to confirm that the peptides were glycosylated, whereas ETD spectra were used for site-localisation of glycosylation sites. All spectra with these modifications were manually annotated.

<u>Click & enrichment of HepG2 secretome:</u>

HepG2-T1$^{-/-}$ and HepG2-T2$^{-/-}$ cells stably transfected with pSBtet-hAGX1$^{3A}$-T1$^{WT}$, pSBtet-hAGX1$^{3A}$-T1$^{DM}$, pSBtet-hAGX1$^{3A}$-T2$^{WT}$, or pSBtet-hAGX1$^{3A}$-T2$^{DM}$ were seeded into one 10 cm dish per treatment, using 8 mL growth medium without hygromycin B. Cells were directly induced with 0.5 µg/mL Dox. After 24 h, cells were Dox-induced again and fed with either caged GalNAc-1-phosphate analog **5** (25 µM) or DMSO vehicle. After another 24 h, the medium was aspirated, and cells were washed with pre-warmed serum-free low-Glc DMEM. Serum-free medium was added, cells were treated with Dox again, and either GalNAc-1-phosphate analog **5** or DMSO vehicle again. Cells were incubated for 20 h. Conditioned supernatant was collected and centrifuged for 5 min at 3000 g. The supernatant was concentrated to 2 mL using an Amicon Ultra-15 centrifuge filter (10 kDa MWCO, Millipore). Samples were stored at -80 °C until further use.

In a repeat experiment, HepG2-T1$^{-/-}$ stably transfected with pSBtet-hAGX1$^{3A}$-T1$^{DM}$ and HepG2-T2$^{-/-}$ cells stably transfected with pSBtet-hAGX1$^{3A}$-T2$^{DM}$ were treated with GalNAc-1-phosphate analog **5** and treated the same way.

Samples were treated with PNGase F (Promega, 5 µL of a 1:10 dilution in PBS) and incubated for 4 h at 37 °C. Samples were then treated in that order with 600 µM BTTAA, 300 µM $CuSO_4$, 2.5 mM sodium ascorbate, 2.5 mM aminoguanidinium chloride, and 50 µM biotin probe **10**.

The click reaction was allowed to proceed for 3 h at r.t. under inversion. The samples were transferred into 15 mL Falcon tubes and treated with 5 mL cold (-20 °C) methanol. Samples were left at -80 °C overnight, when a white precipitate had formed. Samples were centrifuged (3700 g, 20 min, 4 °C), the supernatant was discarded, and pellets were washed with 5 mL methanol twice, with centrifugation each time.

The supernatant was completely removed and the pellet air-dried. Samples were treated with 250 µL 0.1% (w/v) Rapigest. Samples were sonicated in a water bath for 25 min and centrifuged (3700 g, 5 min, 4 °C). The supernatants were transferred to a fresh tube, and pellets were treated with 250 µL 6 M urea in PBS. Samples were sonicated and centrifuged again, the pellets treated with 250 µL PBS, sonicated and centrifuged again. All supernatants from solubilization steps were combined, and samples were diluted with PBS to 2 mL. Protein concentration was determined by BCA.

Beads (UltraLink PLUS Streptavidin, 100 µL slurry = 50 µL settled resin per sample) were washed with PBS twice (harvest 30 s at 1000 g in table top) in low-bind tubes (Eppendorf) and treated with the protein solutions. Samples were incubated for 16 h at room temperature under rotation. The beads were harvested and the supernatant was discarded. The beads were treated as described (*27*). Briefly, beads were washed sequentially with 0.1% RapiGest in PBS (2x), 6 M urea in PBS (3x), and PBS (3x), and resuspended in PBS (200 µL). Beads were treated with 100 mM DTT (Thermo Fisher) in PBS (10 µL), and incubated at r.t. while shaking (950 rpm) for 30 min. Then, 500 mM iodoacetamide (Sigma Aldrich) in PBS (4 µL) was added and samples were shaken for another 30 min in the dark. Beads were harvested and washed with PBS and 50 mM ammonium bicarbonate in LC/MS-grade water (3x, ABC buffer). Beads were resuspended in ABC buffer (200 µL), treated with RapiGest to a final concentration of 0.05% (w/v), and trypsin (0.5 µg in 1 µL ABC buffer). Samples were shaken at 37 °C for 2-3 h, and another 0.5 µg trypsin was added. The reactions were shaken overnight at 37 °C. The beads were harvested, washed with ABC (200 µL) and with LC-MS grade water (3x200 µL). Supernatants were discarded, and the beads were treated with 2% (v/v) formic acid in water (150 µL). Samples were shaken at r.t. for 30 min, and the cleavage step was repeated. The beads were washed with water (100 µL) and 1:1 MeCN/2% aq. FA (100 µL). All supernatants were combined for each sample, centrifuged (18000 g, 5 min, room temperature) and concentrated by Speedvac. Samples were dissolved in 0.1% (v/v) formic acid in water and used for mass spectrometry (see above).

### Synthetic chemistry

Solvents and reagents were of commercial grade. Anhydrous solvents were obtained from a Dry Solvent System. Moisture-sensitive reactions were carried out in heat-dried glassware and under a nitrogen atmosphere. Thin layer chromatography was performed on Kieselgel 60 F254 glass plates pre-coated with silica gel (0.25 mm thickness). Spots were developed with sugar stain (0.1% (v/v) 3-methoxyphenol, 2.5% (v/v) sulfuric acid in EtOH) or ceric ammonium molybdate stain (5% (w/v) ammonium molybdate, 1% (w/v) cerium(II) sulfate and 10% (v/v) sulfuric acid in water) dipping solutions. Flash chromatography was carried out on Fluka Kieselgel 60 (230-400 mesh). Solvents were removed under reduced pressure using a rotary evaporator and high vacuum (1 mbar).

$^{1}H$, $^{13}C$ and 2D NMR spectra were measured with a Varian AS400 spectrometer or a Varian AS600 spectrometer at 25 °C. Chemical shifts (σ) are reported in parts per million (ppm) relative to the respective residual solvent peaks ($CDCl_3$: σ 7.26 in $^{1}H$ and 77.16 in $^{13}C$ NMR; acetone-$D_6$: σ 2.05 in $^{1}H$ and 29.84 in $^{13}C$ NMR). Two-dimensional NMR experiments (HH-COSY, CH-HSQC) were performed to assign peaks in $^{1}H$ spectra. The following abbreviations are used to indicate peak multiplicities: s singlet; d doublet; dd doublet of doublets; dt doublet of triplets; m multiplet. Coupling constants (*J*) are reported in Hertz (Hz). High resolution mass spectrometry by electrospray ionization (ESI-HRMS) was performed at Stanford University Mass Spectrometry, with a micrOTOF-Q II hybrid quadrupole time-of-flight mass spectrometer (Bruker) equipped with an Agilent 1260 UPLC.

Compound **SI-5** was purchased from Oakwood Chemicals (West Columbia, USA).

**Bis(*S*-acetyl-2-thioethyl) 3,4,6-tri-*O*-acetyl-2-deoxy-2-(5-hexynoyl)amido-α-D-galactopyranosyl phosphate (5)**

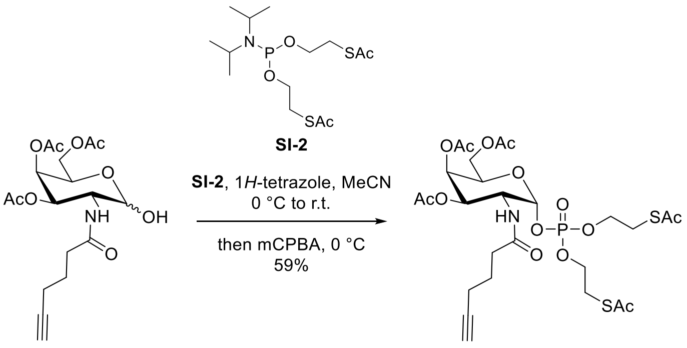

**SI-1** **5**

To a stirred solution of lactol **SI-1** (*8*) (100 mg, 250 umol) in MeCN (0.5 mL) were added at 0 °C phosphoramidite **SI-2** (*14*) (133 mg, 362 umol) in 0.75 mL MeCN and 1*H*-tetrazole (27.1 mg, 388 umol, 873 µL of a 3% (w/v) solution in MeCN). The reaction was warmed to room temperature and stirred for 1 h. Phosphoamidite **SI-2** (34 mg, 93 umol) and 1*H*-tetrazole (235 µL, 104 umol) were added to drive the reaction to completion. The mixture was stirred for 20 min at room temperature, cooled to 0 °C and treated with mCPBA (64.7 mg, 616 umol). The reaction was stirred for 2 h at that temperature and quenched with 10% aq. $Na_2SO_3$ (10 mL). The solution was extracted with $CH_2Cl_2$ (5x10 mL), the combined organic layers were dried over $MgSO_4$, filtered and concentrated. The residue was purified by flash chromatography (hexanes/EtOAc 1:0 to 1:1 with 0.5% (v/v) $NEt_3$) to give phosphotriester **5** (101 mg, 147 umol, 59%) as a clear oil. $R_f$ (hexanes/EtOAc with 0.5% (v/v) $NEt_3$, TLC plates pre-neutralized with hexanes/2% $NEt_3$) = 0.25. $^1$H NMR (600 MHz, $CDCl_3$) δ 6.24 (d, *J* = 9.2 Hz, 1H), 5.75 (dd, *J* = 5.6, 3.3 Hz, 1H), 5.44 (dd, *J* = 3.2, 1.4 Hz, 1H), 5.18 (dd, *J* = 11.5, 3.2 Hz, 1H), 4.81 – 4.60 (m, 1H), 4.40 (td, *J* = 6.6, 1.4 Hz, 1H), 4.29 – 3.93 (m, 4H), 3.34 – 3.06 (m, 6H), 2.37 – 2.30 (m, 8H), 2.28 – 2.17 (m, 2H), 2.15 (s, 3H), 2.03 (s, 3H), 1.99 (s, 3H), 1.95 (t, *J* = 2.6 Hz, 1H), 1.88 – 1.76 (m, 2H); $^{13}$C NMR (150 MHz, $CDCl_3$) δ 195.3, 194.9, 172.8, 170.7, 170.4, 170.2, 97.3, 97.3, 83.4, 69.5, 69.3, 68.8, 67.4, 67.3, 66.9, 66.8, 66.6, 66.5, 66.5, 66.5, 61.5, 47.6, 47.6, 47.5, 47.4, 34.8, 34.7, 30.8, 30.7, 30.6, 29.2, 29.2, 29.2, 24.1, 24.0, 20.8, 20.7, 20.6, 17.8, 17.7; HRMS (ESI) calcd. for $C_{26}H_{38}NO_{14}PS_2Na$ ($M+Na^+$) 706.1369 found 706.1349 *m/z*.

**Bis(*S*-acetyl-2-thioethyl) 3,4,6-tri-*O*-acetyl-2-deoxy-2-(2-(S)-methyl-5-hexynoyl)amido-α-D-galactopyranosyl phosphate (6)**

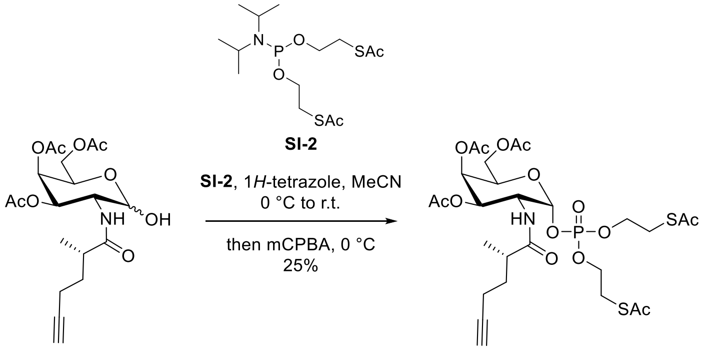

**SI-3** **6**

This reaction is not optimized.

To a stirred solution of lactol **SI-3** (*8*) (30 mg, 73 umol) in MeCN (0.73 mL) were added at 0 °C phosphoramidite **SI-2** (30 mg, 81 umol) in 0.4 mL toluene and 1*H*-tetrazole (9.1 mg, 130 umol, 252 µL of a 3% (w/v) solution in MeCN). The reaction was warmed to room temperature and stirred for 1 h. Phosphoamidite **SI-2** (30 mg, 81 umol) was added to drive the reaction to completion. The mixture was stirred for 20 min at room temperature, cooled to -20 °C and treated with mCPBA (18.6 mg, 108 umol). The reaction was stirred for 2 h at that temperature and quenched with 10% aq. $Na_2SO_3$ (10 mL). The solution was extracted with $CH_2Cl_2$ (5x10 mL), the combined organic layers were dried over $MgSO_4$, filtered and concentrated. The residue was purified by flash chromatography (hexanes/EtOAc 1:0 to 5:1 to 3:1to 1:1 with 0.5% (v/v) $NEt_3$) and size exclusion chromatography (Sephadex LH-20, solvent $CH_2Cl_2$ 4:1) to give phosphotriester **6** (12.5 mg, 18 umol, 25%) as a clear oil. $^1$H NMR (400 MHz, acetone-$D_6$) δ 7.36 (d, *J* = 8.3 Hz, 1H), 5.78 (dd, *J* = 6.0, 3.4 Hz, 1H), 5.50 (dd, *J* = 3.2, 1.4 Hz, 1H), 5.17 (dd, *J* = 11.8, 3.2 Hz, 1H), 4.71 – 4.44 (m, 2H), 4.31 – 4.03 (m, 6H), 3.22 (t, *J* = 6.5 Hz, 4H), 2.62 – 2.44 (m, 1H), 2.44 – 2.28 (m, 7H), 2.19 – 2.11 (m, 5H), 2.01 (s, 3H), 1.94 (s, 3H), 1.90 – 1.75 (m, 1H), 1.61 – 1.49 (m, 1H), 1.12 (d, *J* = 6.9, 1.6 Hz, 3H); $^{13}$C NMR (100 MHz, acetone-$D_6$) δ 195.2, 177.5, 176.5, 170.7, 170.6, 170.4, 97.6, 84.3, 70.2, 69.4, 67.9, 67.7, 66.9, 62.4, 48.1, 40.3, 33.8, 20.7, 20.6, 18.0, 16.8; HRMS (ESI) calcd. for $C_{27}H_{40}NO_{14}PS_2Na$ (M+Na$^+$) 720.1526 found 720.1523 *m/z*.

**Bis(*S*-acetyl-2-thioethyl) 3,4,6-tri-*O*-acetyl-2-deoxy-2-(2-(R)-methyl-5-hexynoyl)amido-α-D-galactopyranosyl phosphate (7)**

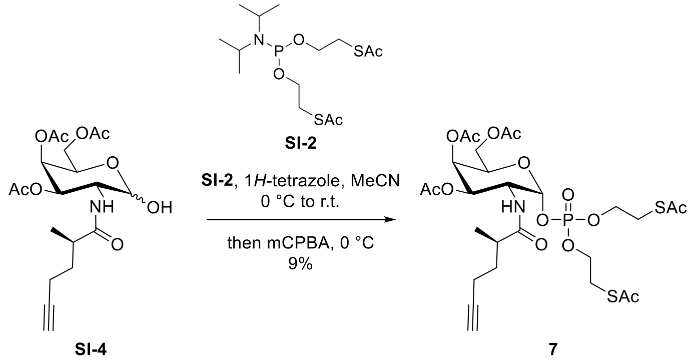


This reaction is not optimized.

To a stirred solution of lactol **SI-4** (*8*) (71 mg, 171 umol) in MeCN (0.73 mL) were added at 0 °C phosphoramidite **SI-2** (75 mg, 202 umol) in 0.4 mL toluene and 1*H*-tetrazole (21.5 mg, 307 umol, 716 µL of a 3% (w/v) solution in MeCN). The reaction was warmed to room temperature and stirred for 1 h. Phosphoamidite **SI-2** (30 mg, 81 umol) was added to drive the reaction to completion. The mixture was stirred for 20 min at room temperature, cooled to -20 °C and treated with mCPBA (50 mg, 290 umol). The reaction was stirred for 2 h at that temperature and quenched with 10% aq. $Na_2SO_3$ (10 mL). The solution was extracted with $CH_2Cl_2$ (5x10 mL), the combined organic layers were dried over $MgSO_4$, filtered and concentrated. The residue was purified by flash chromatography (hexanes/EtOAc 1:0 to 5:1 to 3:1to 1:1 with 0.5% (v/v) $NEt_3$) and size exclusion chromatography (Sephadex LH-20, solvent $CH_2Cl_2$ 4:1) to give phosphotriester **7** (11 mg, 16 µmol, 9%) as a clear oil. $^1$H NMR (600 MHz, acetone-$D_6$) δ 7.35 (d, *J* = 8.2 Hz, 1H), 5.81 (m, 1H), 5.50 (s, 1H), 5.18 (m, 1H), 4.59 – 4.43 (m, 2H), 4.33 – 4.01 (m, 6H), 3.31 – 3.07 (m, 4H), 2.55 – 2.45 (m, 1H), 2.40 – 2.30 (m, 7H), 2.25 – 2.12 (m, 5H), 2.01 (s, 3H), 1.97 – 1.85 (m, 4H), 1.57 – 1.48 (m, 1H), 1.09 (d, *J* = 7.0 Hz, 3H); $^{13}$C NMR (150 MHz, acetone-$D_6$) δ 195.2, 176.6, 170.6, 170.4, 97.6, 97.6, 84.7, 70.1, 69.5, 68.0, 67.8, 67.1, 66.9, 62.4, 48.3, 40.4, 33.4, 30.6, 30.6, 20.3, 20.6, 18.3, 16.9; HRMS (ESI) calcd. for $C_{27}H_{40}NO_{14}PS_2Na$ (M+Na$^+$) 720.1526 found 720.1515 *m/z*.

**3-Hydroxypropyl (6-azidomethyl)nicotinate (SI-7)**

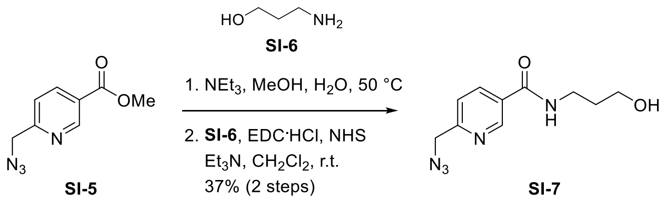


To a stirred solution of ester **SI-5** (58 mg, 0.30 mmol) in 5:2 MeOH/water (1.5 mL) was added at room temperature triethylamine (0.3 mL). The reaction was warmed to 50 °C and stirred for 20 h at that temperature, when the starting material was converted to a lower-running spot (TLC $CH_2Cl_2$/MeOH 10:1). The solvents were evaporated, and the residue was co-evaporated with toluene (3x5 mL) to give the intermediate triethylammonium salt as a yellow oil.

To a stirred solution of the intermediate triethylammonium salt in $CH_2Cl_2$ (1.5 mL) were added at room temperature triethylamine (164 µL, 1.18 umol), EDC hydrochloride (116 mg, 0.6 mmol) and *N*-hydroxysuccinimide (52 mg, 0.45 mmol). The mixture was stirred for 2 h, and 3-amino-1-propanol **SI-6** (72 mg, 0.95 mmol) was added. The reaction was stirred for 20 h, diluted with $CH_2Cl_2$ (5 mL), and filtered through cotton wool. The solution was concentrated, and the residue was purified by flash chromatography ($CH_2Cl_2$/MeOH 99:1 to 95:5) to give amide **SI-7** (26 mg, 111 umol, 37% over two steps) as a clear oil. $R_f$ ($CH_2Cl_2$/MeOH 20:1) = 0.3. $^1$H NMR (400 MHz, $CDCl_3$) δ 9.28 (d, *J* = 2.3 Hz, 1H), 8.49 (dd, *J* = 8.1, 2.3 Hz, 1H), 7.77 (d, *J* = 8.1 Hz, 1H), 7.44 (br s, 1H), 4.88 (s, 2H), 4.13 (t, *J* = 5.5 Hz, 2H), 3.99 (q, *J* = 5.9 Hz, 2H), 2.70 (br s, 1H), 2.25 – 2.11 (m, 2H); $^{13}$C NMR (100 MHz, $CDCl_3$) δ 165.9, 158.8, 147.9, 136.4, 129.4, 121.8, 60.8, 55.4, 38.2, 31.7. HRMS (ESI) calcd. for $C_{10}H_{13}N_5O_2Na$ (M+Na$^+$) 258.0966 found 258.0968 *m/z*.

**1,3,4,6-Tri-*O*-acetyl-2-deoxy-2-(5-hexynoyl)amido-αβ-D-glucopyranoside (9)**

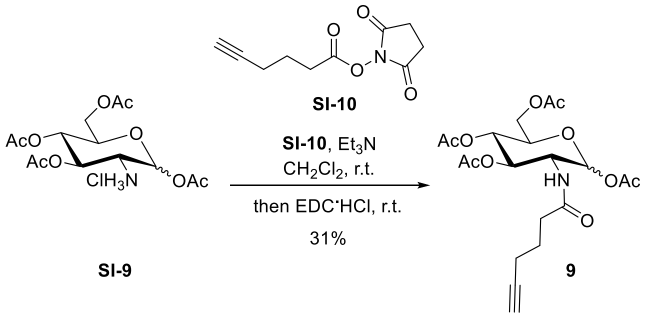


To a stirred solution of ammonium salt **SI-10** (200 mg, 0.52 mmol) in 5:2 $CH_2Cl_2$ (2.6 mL) were added at room temperature active ester **SI-9** (130 mg, 0.63 mmol) and triethylamine (145 µL, 1.04 mmol). The reaction was stirred for 16 h and treated with EDC hydrochloride (99.8 mg, 0.52 mmol). The mixture was stirred for 48 h, the reaction was diluted with EtOAc (10 mL), washed with 1 N HCl (2x5 mL), sat. aq. $NaHCO_3$ (5 mL) and brine (5 mL). The combined organic layers were dried over $MgSO_4$, filtered and concentrated. The residue was purified by flash chromatography (hexanes/EtOAc 2:1 to 1:2 to 1:4 to give amide **9** (72 mg, 0.16 mmol, 31%) as a clear oil. $^1$H NMR (400 MHz, $CDCl_3$) δ 5.88 (d, *J* = 10.2 Hz, 1H), 5.69 (d, *J* = 8.7 Hz, 1H), 5.30 – 5.01 (m, 2H), 4.43 – 4.20 (m, 2H), 4.11 (dd, *J* = 12.5, 2.2 Hz, 1H), 3.83 (ddd, *J* = 9.9, 4.7, 2.2 Hz, 1H), 2.32 – 2.15 (m, 4H), 2.10 (s, 3H), 2.07 (s, 3H), 2.03 (d, *J* = 1.5 Hz, 6H), 1.95 (t, *J* = 2.6 Hz, 1H), 1.77 (p, *J* = 7.0 Hz, 2H). $^{13}$C NMR (100 MHz, $CDCl_3$) δ 172.4, 171.3, 170.8, 169.6, 169.5, 92.6, 83.2, 72.9, 72.6, 69.6, 68.1, 61.8, 52.8, 34.9, 24.0, 21.0, 20.8, 20.8, 20.7, 17.6. HRMS (ESI) calcd. for $C_{20}H_{27}NO_{10}Na$ (M+Na$^+$) 464.1532 found 464.1513 *m/z*.

**Silane probe 10**

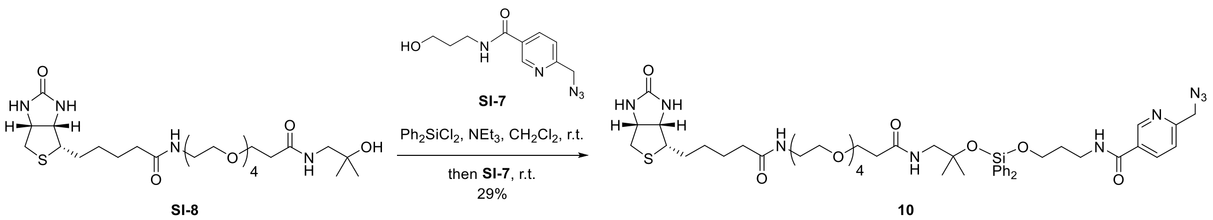

To a stirred solution of alcohol **SI-8** (10 mg, 17.7 umol) in $CH_2Cl_2$ (0.5 mL) were added at room temperature triethylamine (50 µL, 360 umol) and dichlorodiphenylsilane (22.4 mg, 88.5 umol). The mixture was stirred for 3 h, and alcohol **SI-7** (36 mg,153 umol in 0.2 mL $CH_2Cl_2$) was added. The reaction was stirred for 20 h, quenched with sat. aq. $NaHCO_3$ (3 mL) and diluted with $CH_2Cl_2$ (5 mL). The layers were separated, and the aqueous phase was extracted with $CH_2Cl_2$ (5x5 mL). The combined organic layers were dried over $Na_2SO_4$, filtered and concentrated. The residue was purified by flash chromatography ($CH_2Cl_2$/MeOH 1:0 to 95:5 with 1% $NEt_3$, then EtOAc/MeOH 1:0 to 9:1 with 1% $NEt_3$) and size exclusion chromatography (Sephadex LH-20, $CH_2Cl_2$/MeOH 1:1) to give diphenyldisiloxane **10** (5 mg, 5.1 umol, 29%) as a clear oil. $R_f$ ($CH_2Cl_2$/MeOH 10:1 with 1% $NEt_3$) = 0.55. $^1$H NMR (600 MHz, $CDCl_3$) δ 8.93 (d, *J* = 2.2 Hz, 1H), 8.05 (dd, *J* = 8.1, 2.3 Hz, 1H), 7.66 – 7.59 (m, 4H), 7.44 – 7.40 (m, 2H), 7.39 – 7.33 (m, 4H), 7.28 (d, *J* = 7.5 Hz, 1H), 7.24 – 7.20 (m, 1H), 6.68 (s, 1H), 6.58 (s, 1H), 5.55 (s, 1H), 4.68 (s, 1H), 4.52 (s, 2H), 4.49 – 4.43 (m, 1H), 4.32 – 4.25 (m, 1H), 3.90 (t, *J* = 5.8 Hz, 2H), 3.70 (t, *J* = 6.1 Hz, 2H), 3.61 – 3.52 (m, 16H), 3.43 – 3.38 (m, 2H), 3.36 (d, *J* = 6.1 Hz, 2H), 3.13 (td, *J* = 7.3, 4.5 Hz, 1H), 2.90 (dd, *J* = 12.8, 5.0 Hz, 1H), 2.69 (d, *J* = 12.7 Hz, 1H), 2.43 (t, *J* = 6.0 Hz, 2H), 2.20 (td, *J* = 7.3, 3.1 Hz, 2H), 1.99 – 1.90 (m, 2H), 1.78 – 1.67 (m, 4H), 1.49 – 1.41 (m, 2H), 1.25 (s, 6H). HRMS (ESI) calcd. for $C_{47}H_{67}N_9O_{10}SSiNa$ (M+Na$^+$) 1000.4399 found 1000.4363 *m/z*.

### Spectral characterization

$^1H$ NMR, $CDCl_3$, 600 MHz

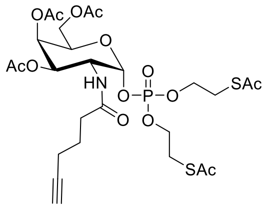


**5**

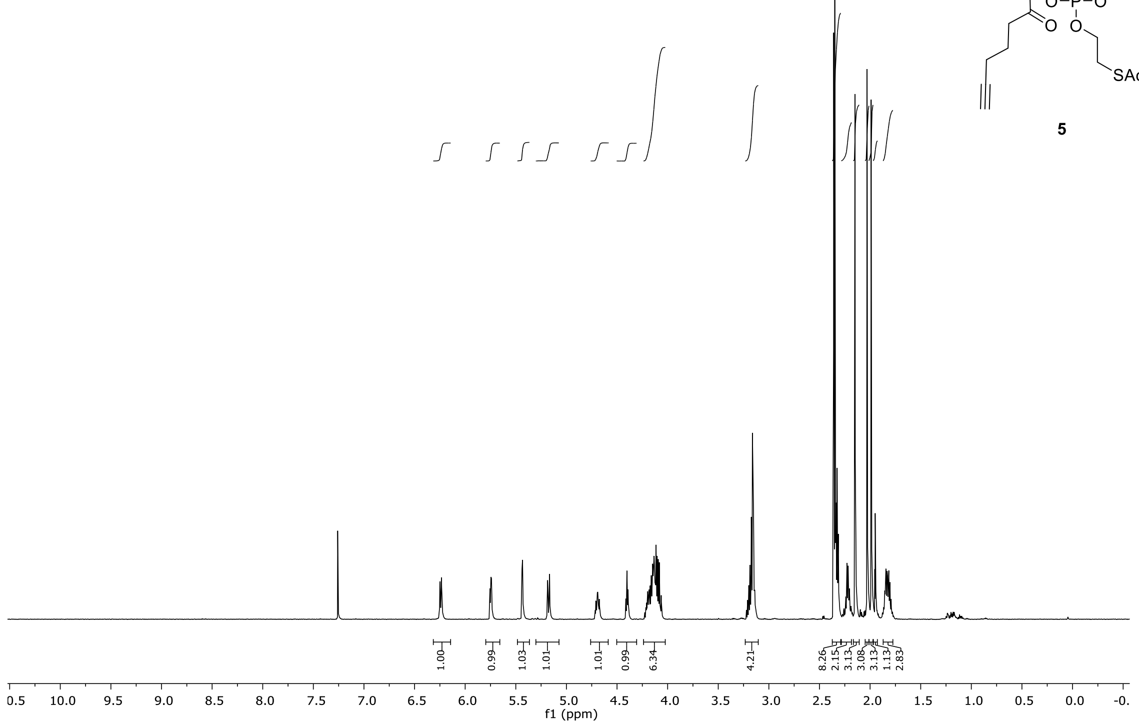


$^{13}C$ NMR, $CDCl_3$, 150 MHz

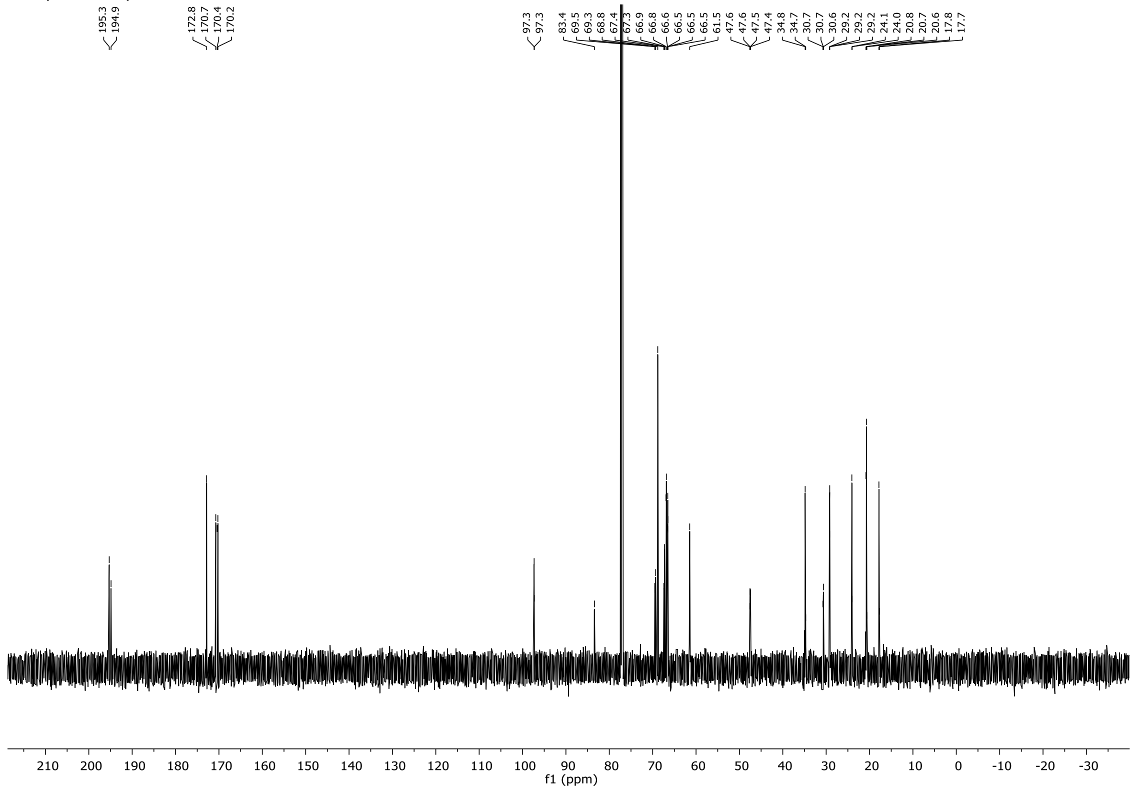


5

$^1$H NMR, acetone-D$_6$, 400 MHz

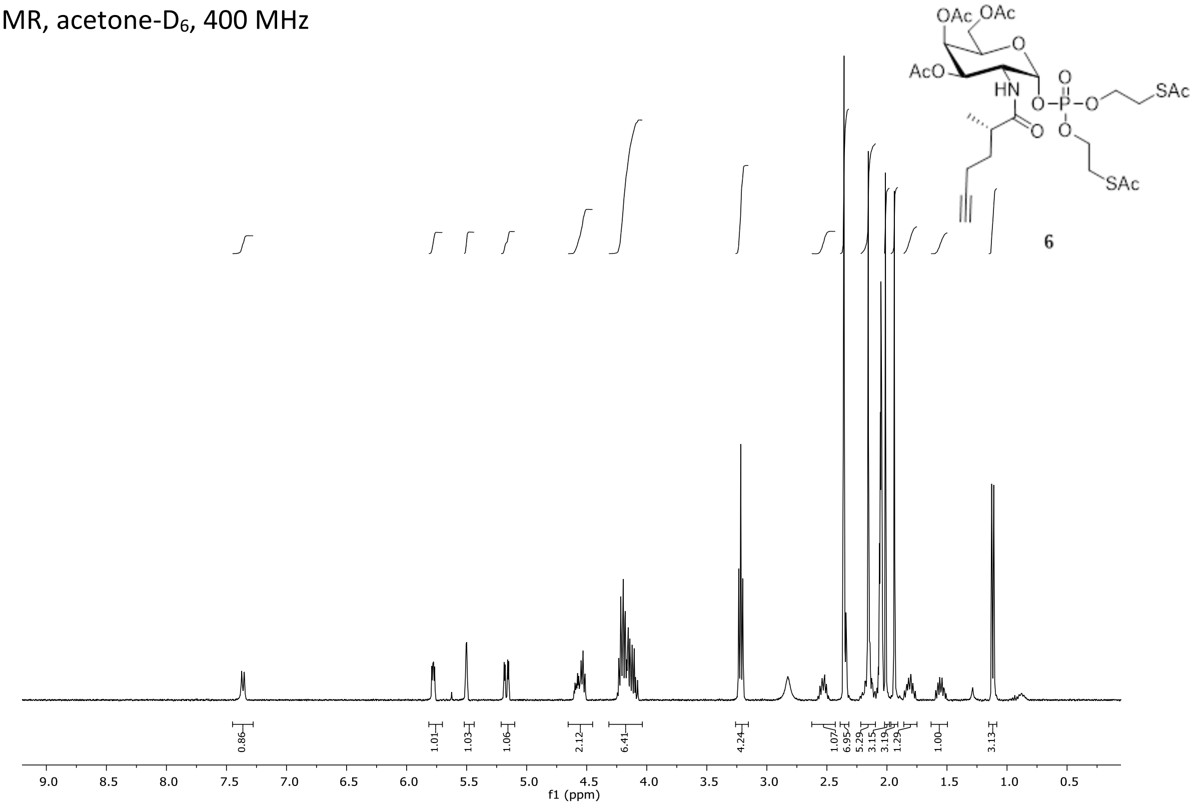


$^{13}$C NMR, acetone-D$_6$, 100 MHz

5

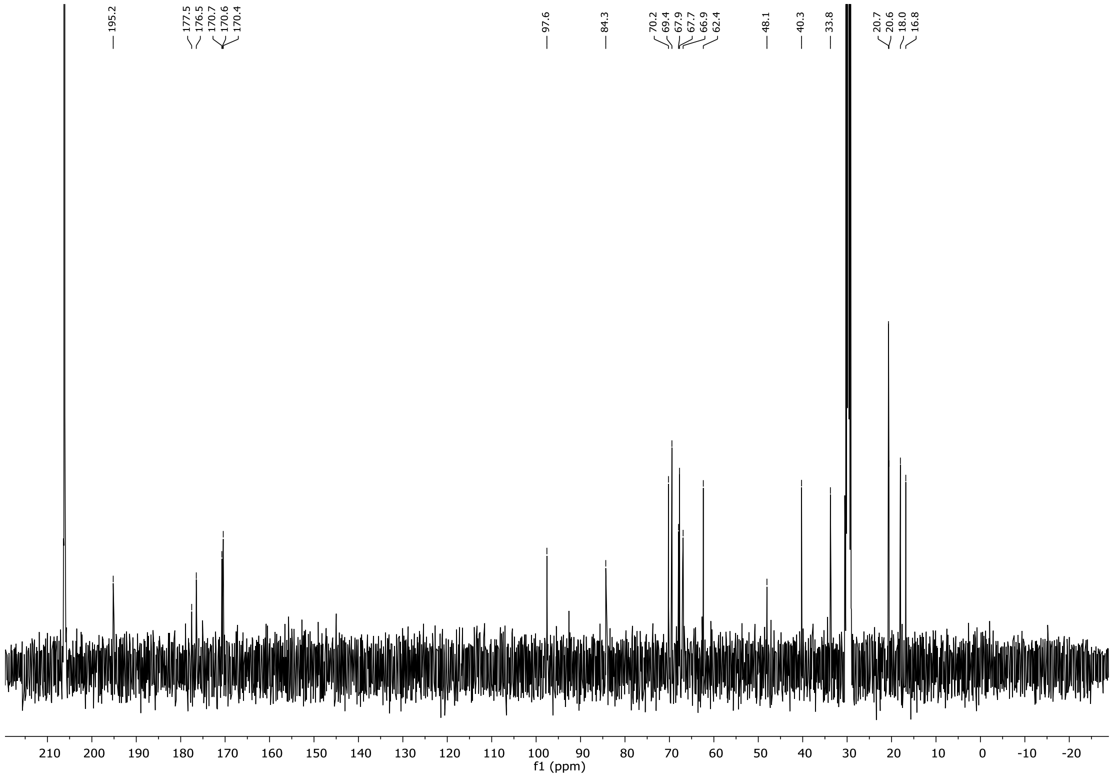

$^{1}H$ NMR, acetone-$D_6$, 400 MHz

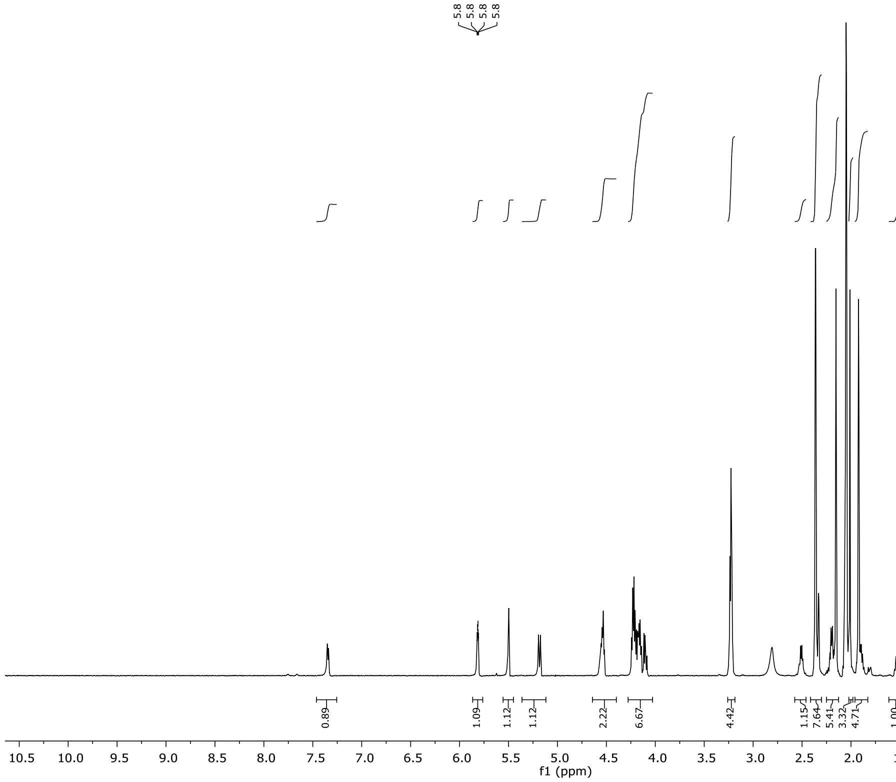


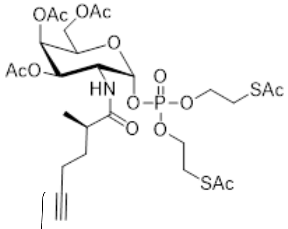


7

$^{13}C$ NMR, acetone-$D_6$, 100 MHz

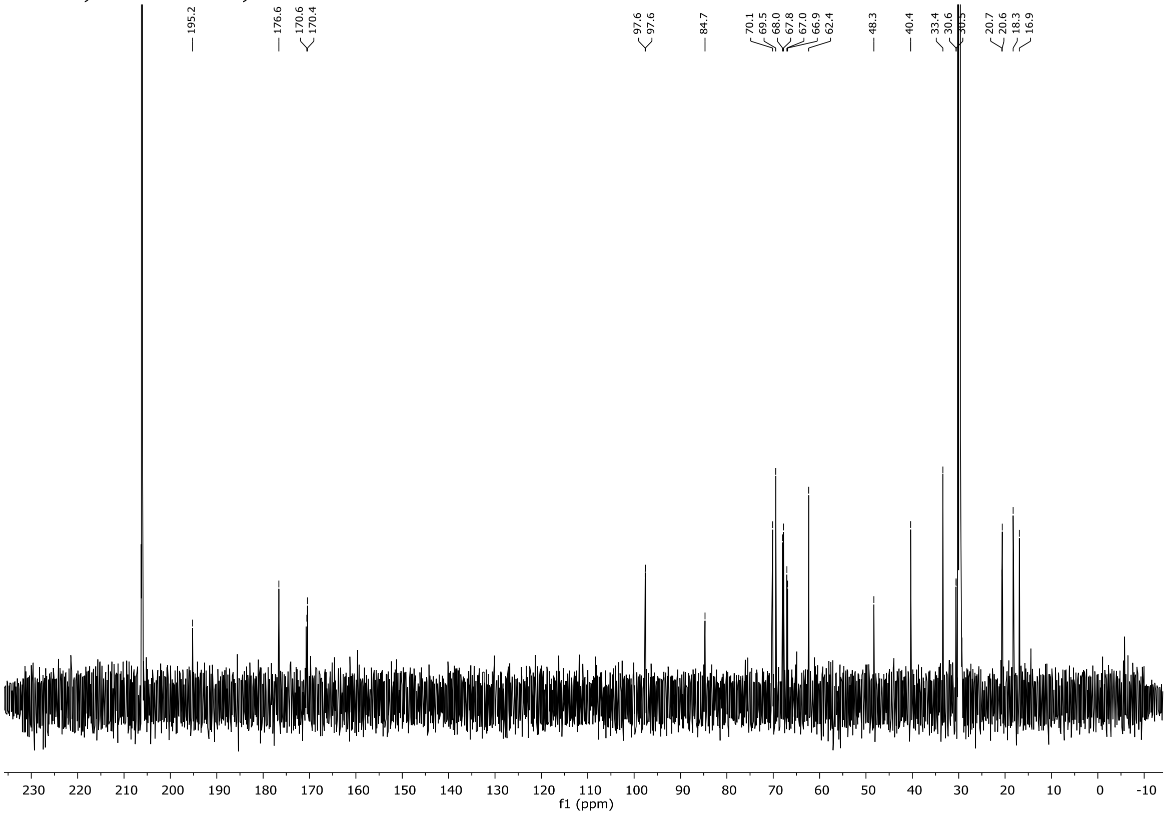

$^{1}H$ NMR, $CDCl_3$, 400 MHz

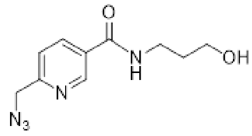


SI-7

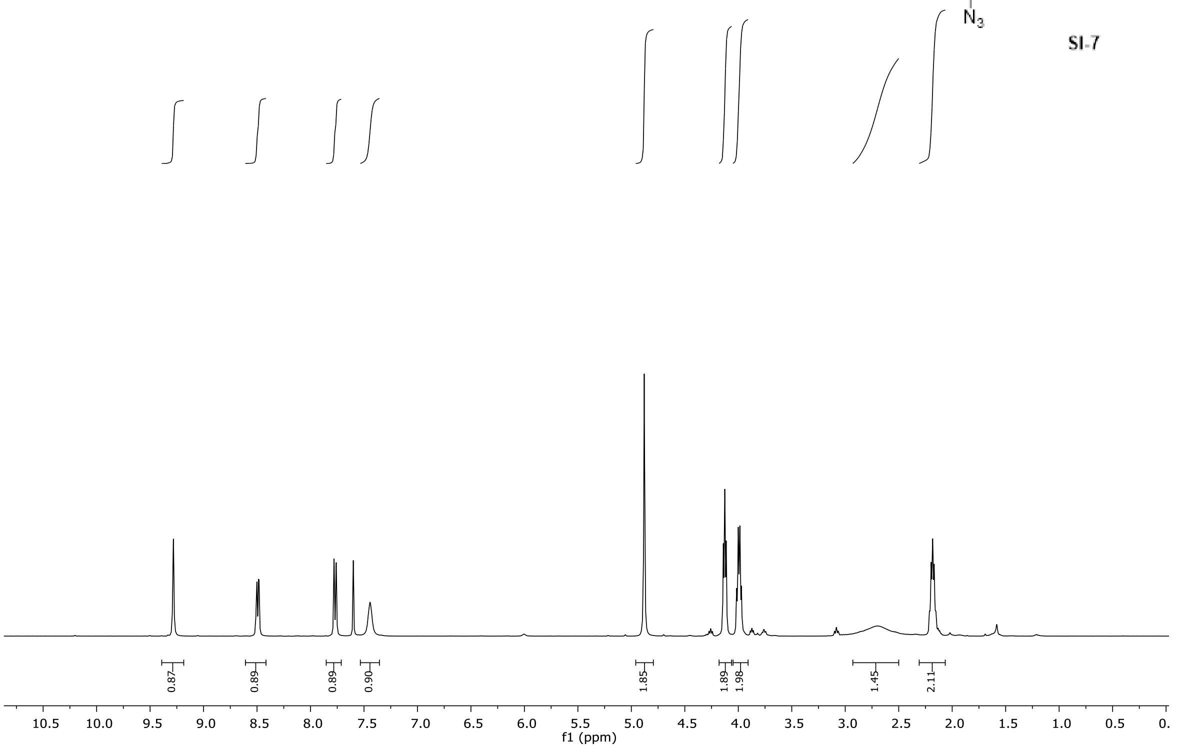


$^{1}H$ NMR, $CDCl_3$, 100 MHz

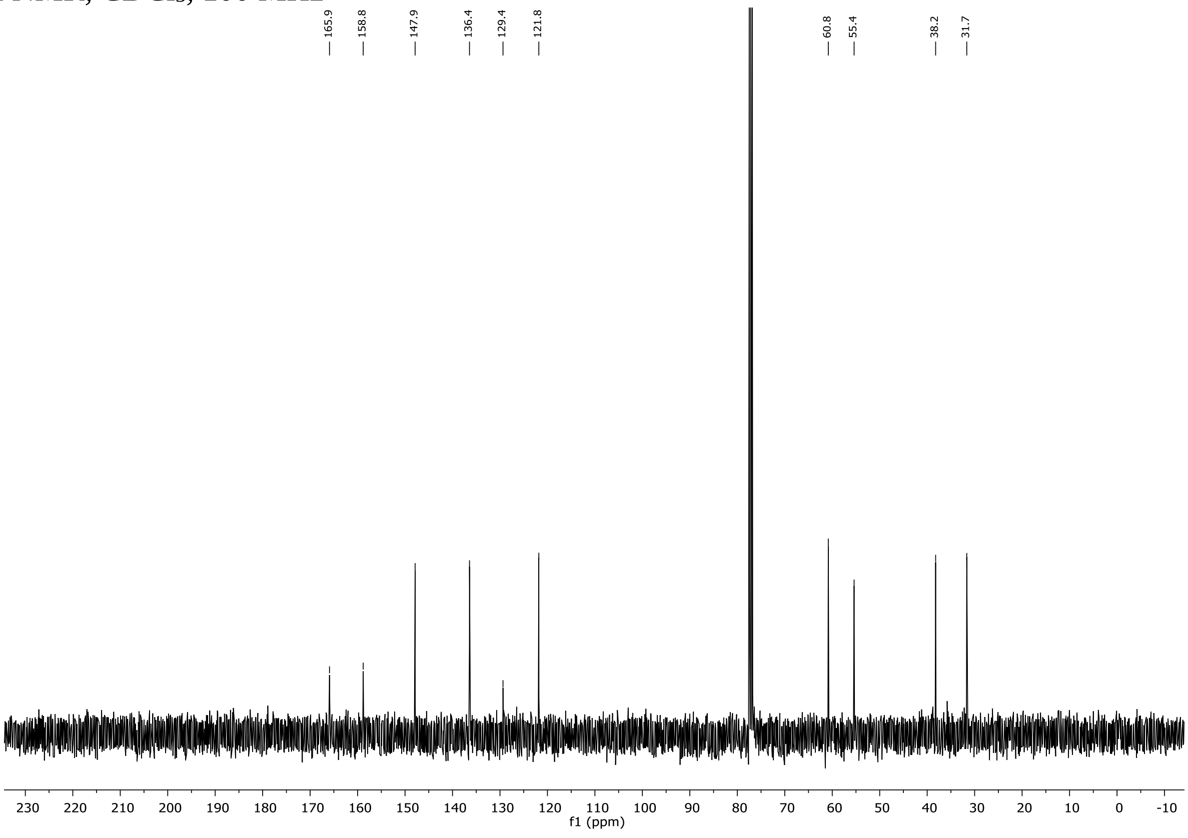

$^1H$ NMR, $CDCl_3$, 400 MHz

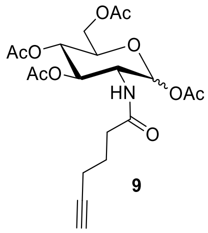


**9**

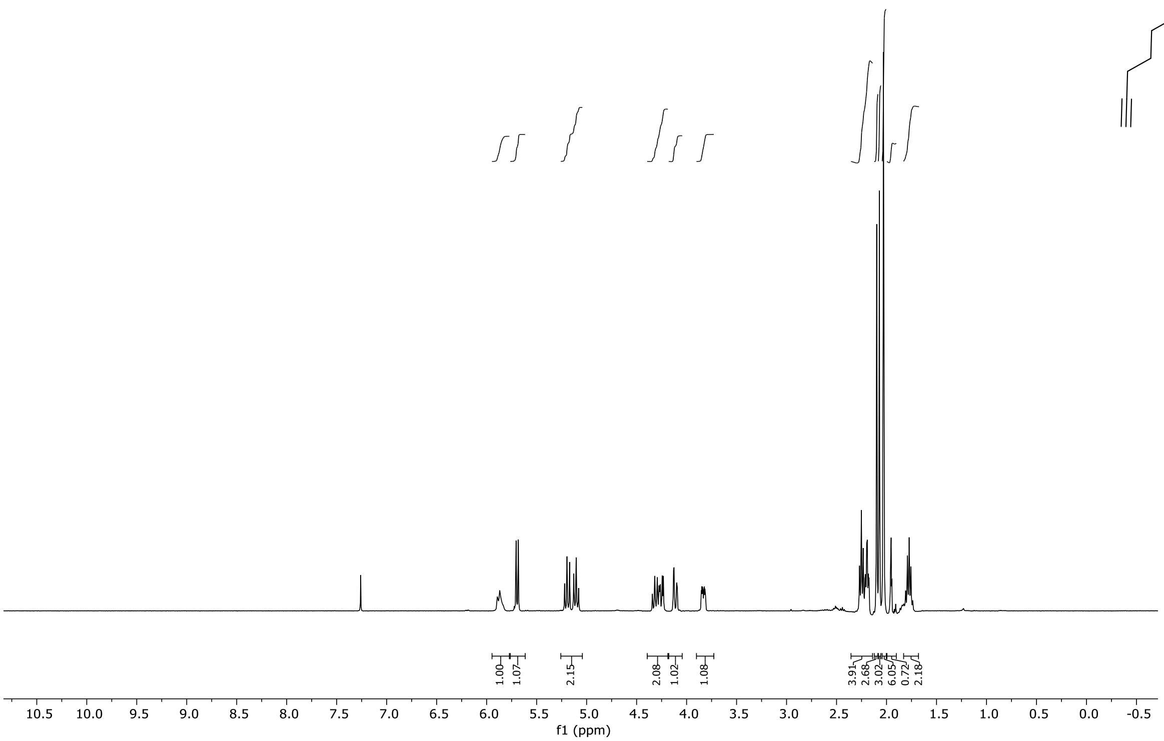


$^1H$ NMR, $CDCl_3$, 600 MHz

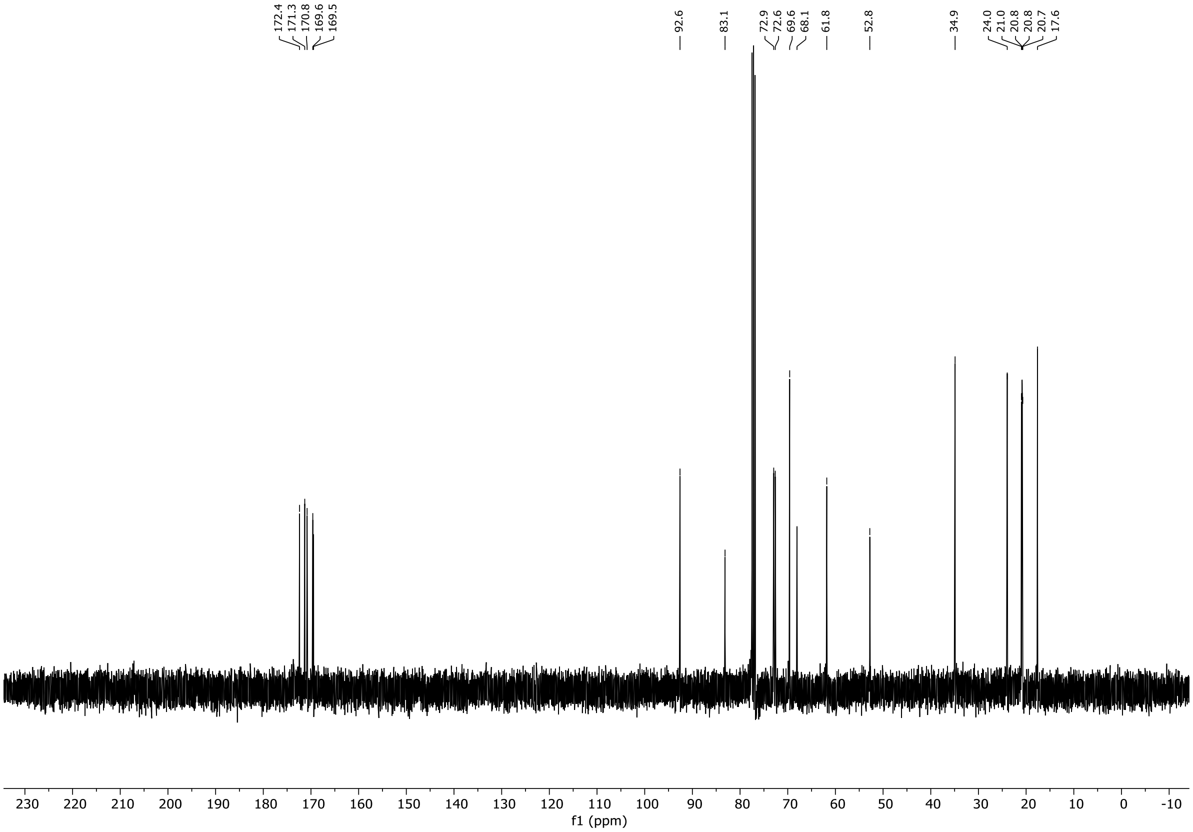

$^1H$ NMR, $CDCl_3$, 600 MHz

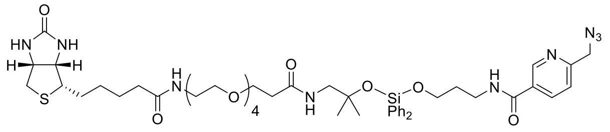


10

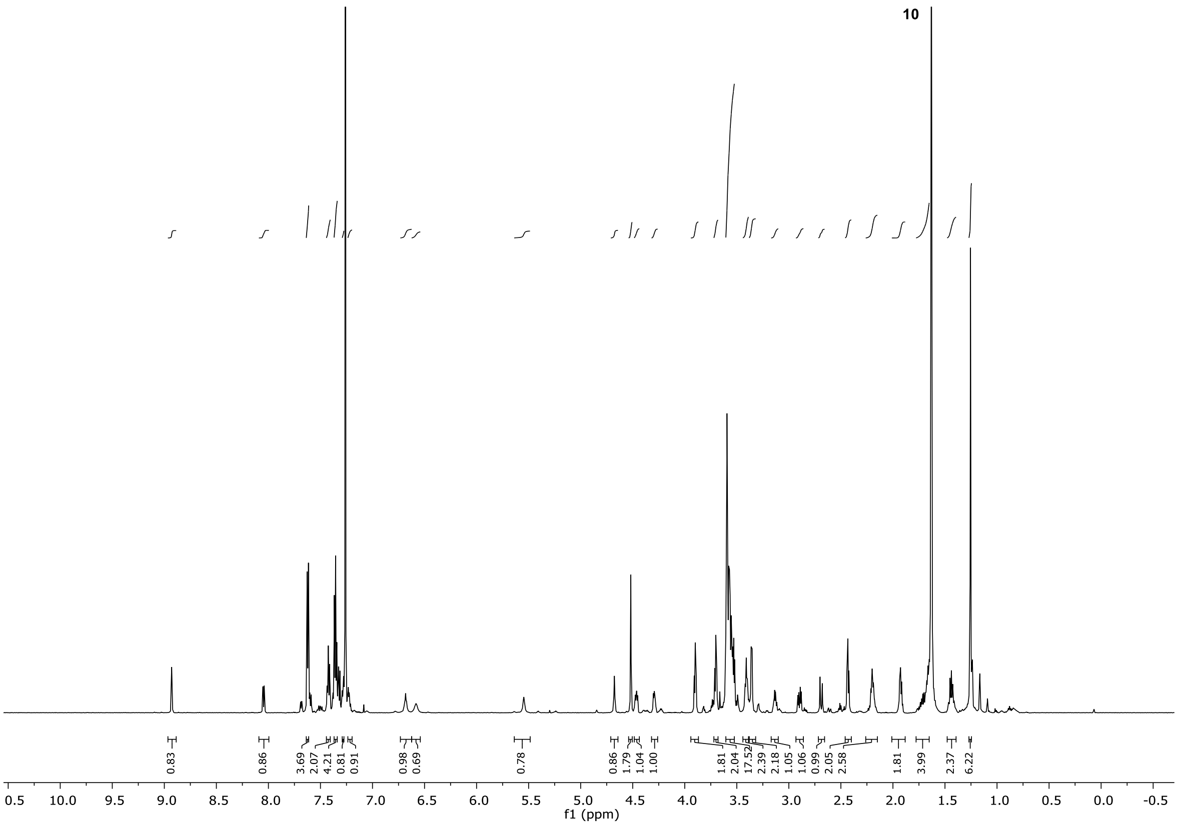
